# The *in vitro* antiviral activity of the anti-hepatitis C virus (HCV) drugs daclatasvir and sofosbuvir against SARS-CoV-2

**DOI:** 10.1101/2020.06.15.153411

**Authors:** Carolina Q. Sacramento, Natalia Fintelman-Rodrigues, Jairo R. Temerozo, Aline de Paula Dias Da Silva, Suelen da Silva Gomes Dias, Carine dos Santos da Silva, André C. Ferreira, Mayara Mattos, Camila R. R. Pão, Caroline S. de Freitas, Vinicius Cardoso Soares, Lucas Villas Bôas Hoelz, Tácio Vinício Amorim Fernandes, Frederico Silva Castelo Branco, Mônica Macedo Bastos, Núbia Boechat, Felipe B. Saraiva, Marcelo Alves Ferreira, Rajith K. R. Rajoli, Carolina S. G. Pedrosa, Gabriela Vitória, Letícia R. Q. Souza, Livia Goto-Silva, Marilia Zaluar Guimarães, Stevens K. Rehen, Andrew Owen, Fernando A. Bozza, Dumith Chequer Bou-Habib, Patrícia T. Bozza, Thiago Moreno L. Souza

**Author notes:** These authors contributed equally to this work (Carolina Q. Sacramento, Natalia Fintelman-Rodrigues). **Correspondence footnote:** Thiago Moreno L. Souza, PhD, Fundação Oswaldo Cruz (Fiocruz), Centro de Desenvolvimento Tecnológico em Saúde (CDTS), Instituto Oswaldo Cruz (IOC), Pavilhão Osório de Almeida, sala 16, Av. Brasil 4365, Manguinhos, Rio de Janeiro - RJ, Brasil, CEP 21060340.

## Abstract

Current approaches of drugs repurposing against 2019 coronavirus disease (COVID-19) have not proven overwhelmingly successful and the severe acute respiratory syndrome coronavirus 2 (SARS-CoV-2) pandemic continues to cause major global mortality. Daclatasvir (DCV) and sofosbuvir (SFV) are clinically approved against hepatitis C virus (HCV), with satisfactory safety profile. DCV and SFV target the HCV enzymes NS5A and NS5B, respectively. NS5A is endowed with pleotropic activities, which overlap with several proteins from SARS-CoV-2. HCV NS5B and SARS-CoV-2 nsp12 are RNA polymerases that share homology in the nucleotide uptake channel. We thus tested whether SARS-COV-2 would be susceptible these anti-HCV drugs. DCV consistently inhibited the production of infectious SARS-CoV-2 in Vero cells, in the hepatoma cell line (HuH-7) and in type II pneumocytes (Calu-3), with potencies of 0.8, 0.6 and 1.1 μM, respectively. Although less potent than DCV, SFV and its nucleoside metabolite inhibited replication in Calu-3 cells. Moreover, SFV/DCV combination (1:0.15 ratio) inhibited SARS-CoV-2 with EC_50_ of 0.7:0.1 μM in Calu-3 cells. SFV and DCV prevented virus-induced neuronal apoptosis and release of cytokine storm-related inflammatory mediators, respectively. Both drugs inhibited independent events during RNA synthesis and this was particularly the case for DCV, which also targeted secondary RNA structures in the SARS-CoV-2 genome. Concentrations required for partial DCV *in vitro* activity are achieved in plasma at Cmax after administration of the approved dose to humans. Doses higher than those approved may ultimately be required, but these data provide a basis to further explore these agents as COVID-19 antiviral candidates.

## 1) Introduction

In these two decades of the 21^st^ century, life-threatening public health emergencies were related to highly pathogenic coronaviruses (CoV), such as severe acute respiratory syndrome (SARS-CoV) in 2002, middle-east respiratory syndrome (MERS-CoV) in 2014[1] and SARS-CoV-2 contemporaneously. After 8 months of the 2019 CoV disease (COVID-19) outbreak, 15 million cases and over 750 thousand deaths were confirmed[2].

To specifically combat COVID-19, the World Health Organization (WHO) launched the global Solidarity trial, initially composed of lopinavir (LPV)/ritonavir (RTV), combined or not with interferon-β (IFN-β), chloroquine (CQ) and remdesivir (RDV) [3]. Lack of clinical benefit paused the enthusiasm for CQ, its analogue hydroxychloroquine and LPV/RTV against COVID-19[4–6]. RDV showed promising results in non-human primates and clinical studies during early intervention[5,7,8]. Nevertheless, RDV’s access may be limited due to its price, and the necessity of intravenous use makes early intervention impracticable and complicates feasibility within many healthcare settings.

Direct-acting antivirals (DDA) against hepatitis C virus (HCV) are among the safest antiviral agents, since they become routinely used in the last five years[9]. Due to their recent incorporation among therapeutic agents, drugs like daclatasvir (DCV) and sofosbuvir (SFV) have not been systematically tested against SARS-CoV or MERS-CoV.

DCV inhibits HCV replication by binding to the N-terminus of non-structural protein (NS5A), affecting both viral RNA replication and virion assembly[10]. NS5A is a multifunctional protein in the HCV replicative cycle, involved with recruitment of host cellular lipid droplets, RNA binding and replication, protein-phosphorylation, cell signaling and antagonism of interferon pathways[10]. In large positive sense RNA viruses, such as SARS-CoV-2, these activities are executed by various viral proteins, especially the non-structural proteins (nsp) 1 to 14[11]. SFV inhibits the HCV protein NS5B, its RNA polymerase[12]. This drug has been associated with antiviral activity against other positive sense RNA viruses, such as Zika (ZIKV), yellow fever (YFV) and chikungunya (CHIKV) viruses [13–16]. With respect to HCV, SFV appears to have a high barrier to the development of resistance. SFV is 2`Me-F uridine monophosphate nucleotide[12]. Hydrophobic protections in its phosphate allow SFV to enter the cells, and then this pro-drug must become the active triphosphorylated nucleotide. Although the cellular enzymes cathepsin A (CatA), carboxylesterase 1 (CES1) and histidine triad nucleotide-binding protein 1 (Hint1) involved with removal of monophosphate protections are classically associated with the hepatic expression[17], they are also present in other tissue, such as the respiratory tract [18–20]. Moreover, the similarities between the SARS-CoV-2 and HCV RNA polymerase provide a rational for studying sofosbuvir as an antiviral for COVID-19 [21]. Using enzymatic assays, sofosbuvir was shown to act as a competitive inhibitor and a chain terminator for SARS-CoV-2 RNA polymerase[22,23]. In human brain organoids, SFV protected neural cells from SARS-CoV-2-induced cell death [24].

Taken collectively, current data provided a bases to investigate whether DCV and SFV could inhibit the production of infectious SARS-CoV-2 particles in physiologically relevant cells. DCV consistently inhibited the production of infectious SARS-CoV-2 in different cells, impairing virus RNA synthesis with an apparently novel mechanism of action, by targeting double-stranded viral RNA. DCV also prevented the release of the inflammatory mediators IL-6 and TNF-α, which are associated with COVID-19 cytokine storm, in SARS-CoV-2-infected primary human monocytes. SFV, which was inactive in Vero cells, inhibited SARS-CoV-2 replication more potently in hepatoma than in respiratory cells. Furthermore, SFV potency appeared to be augmented in the presence of sub-inhibitory concentrations of DCV. These data support further investigation of DCV/SFV for COVID-19. Of interest, concentrations providing sub-maximal inhibition of SARS-CoV-2 by DCV are achieved in plasma at maximal concentration (Cmax) after administration of its approved dose of 60mg once daily, which has considerable scope for dose escalation.

## 2) Results

### 2.1) DCV is more potent than SFV to inhibit the production of infectious SARS-CoV-2 particles

SARS-CoV-2 may infect cell lineages from different organs, but permissive production of infectious virus particles varies according to the cellular systems. Since we wanted to diminish infectious virus titers with studied antiviral drugs, we first compared cell types used in SARS-CoV-2 research with respect to their permissiveness to this virus. Whereas African green monkey kidney cell (Vero E6), human hepatoma (HuH-7) and type II pneumocytes (Calu-3) produced infectious SARS-CoV-2 titers and quantifiable RNA levels (Figure S1), A549 pneumocytes and induced pluripotent human neural stem cells (NSC) displayed limited ability to generate virus progeny, as measured by plaque forming units (PFU) of virus bellow the limit of detection (Figure S1A).

Next, the phenotypic experiments were performed at MOI of 0.01 for Vero cells 24h after infection, and 0.1 for HuH-7 and Calu-3 cells at 48h after infection. Cultures were treated after 1h infection period and cell culture supernatant fractions were harvested to measure infectious SARS-CoV-2 by plaque forming units (PFUs) in Vero cells. DCV consistently inhibited the production of SARS-CoV-2 infectious virus titers in a dose-dependent manner in the all tested cell types (Figure 1), being similarly potent in Vero, HuH-7 and Calu-3 cells, with EC_50_ values ranging between 0.6 to 1.1 μM, without statistical distinction (Table 1). DCV showed limited antiviral activity when viral RNA copies/mL in the culture supernatant fraction (Figures S2) was utilized, suggesting a mechanism unrelated to RNA production.

**Figure 1.**
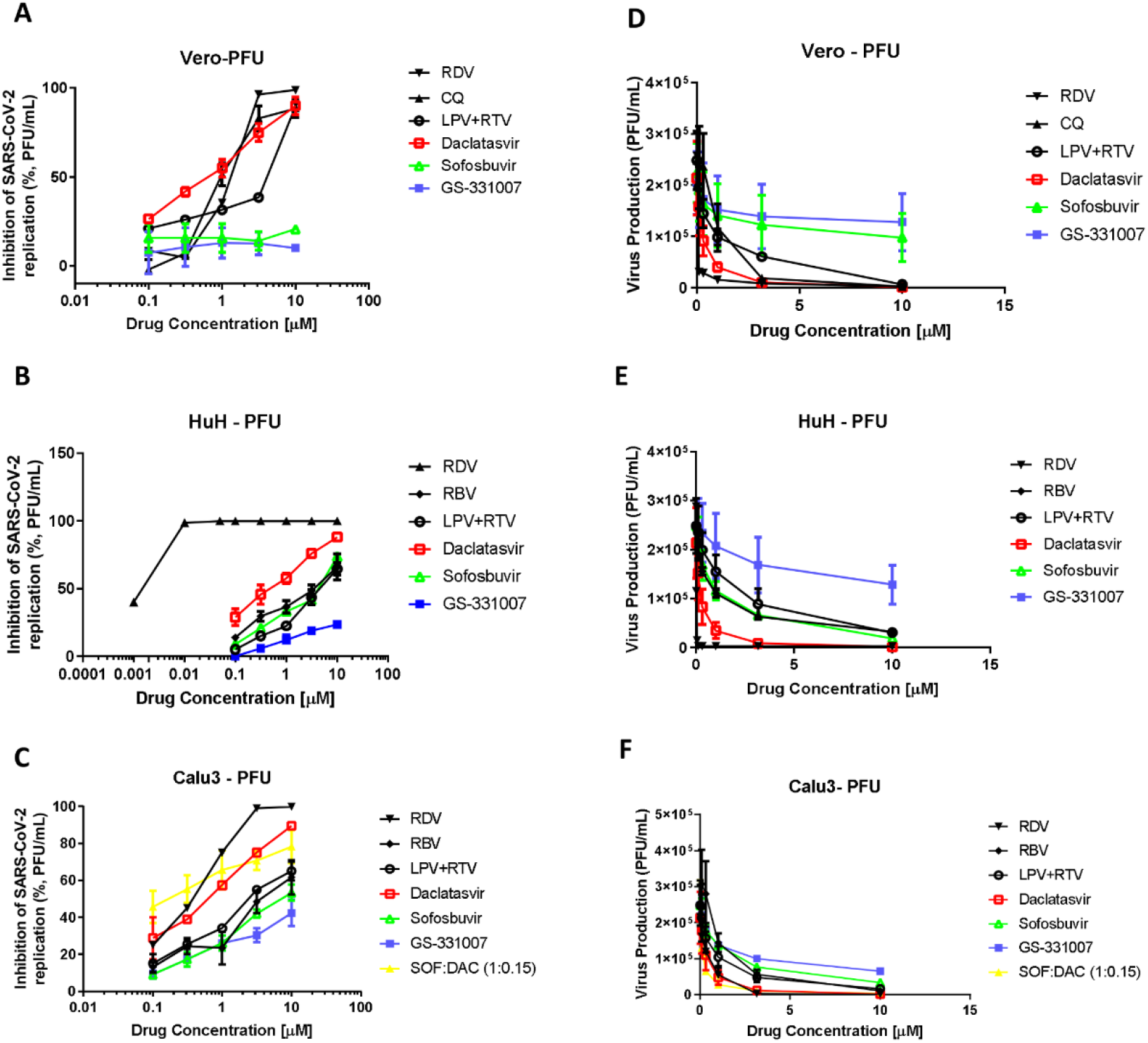
The antiviral activity of daclatasvir (DCV) and sofosbuvir (SFV) against SARS-CoV-2. Vero (A and D), HuH-7 (B and E) or Calu-3 (C and F) cells, at density of 5 × 10^5^ cells/well in 48-well plates, were infected with SARS-CoV-2, for 1h at 37 °C. Inoculum was removed, cells were washed and incubated with fresh DMEM containing 2% fetal bovine serum (FBS) and the indicated concentrations of the DCV, SFV, chloroquine (CQ), lopinavir/ritonavir (LPV+RTV) or ribavirin (RBV). Vero (A and D) were infected with MOI of 0.01 and supernatants were accessed after 24 h. HuH-7 (B and E) and Calu-3 (C and F) cells were infected with MOI of 0.1 and supernatants were accessed after 48 h. Viral replication in the culture supernatant was measured by PFU/mL. Results are displayed as percentage of inhibition (A-C) or virus titers (D-F). The data represent means ± SEM of three independent experiments.

**Table 1.**
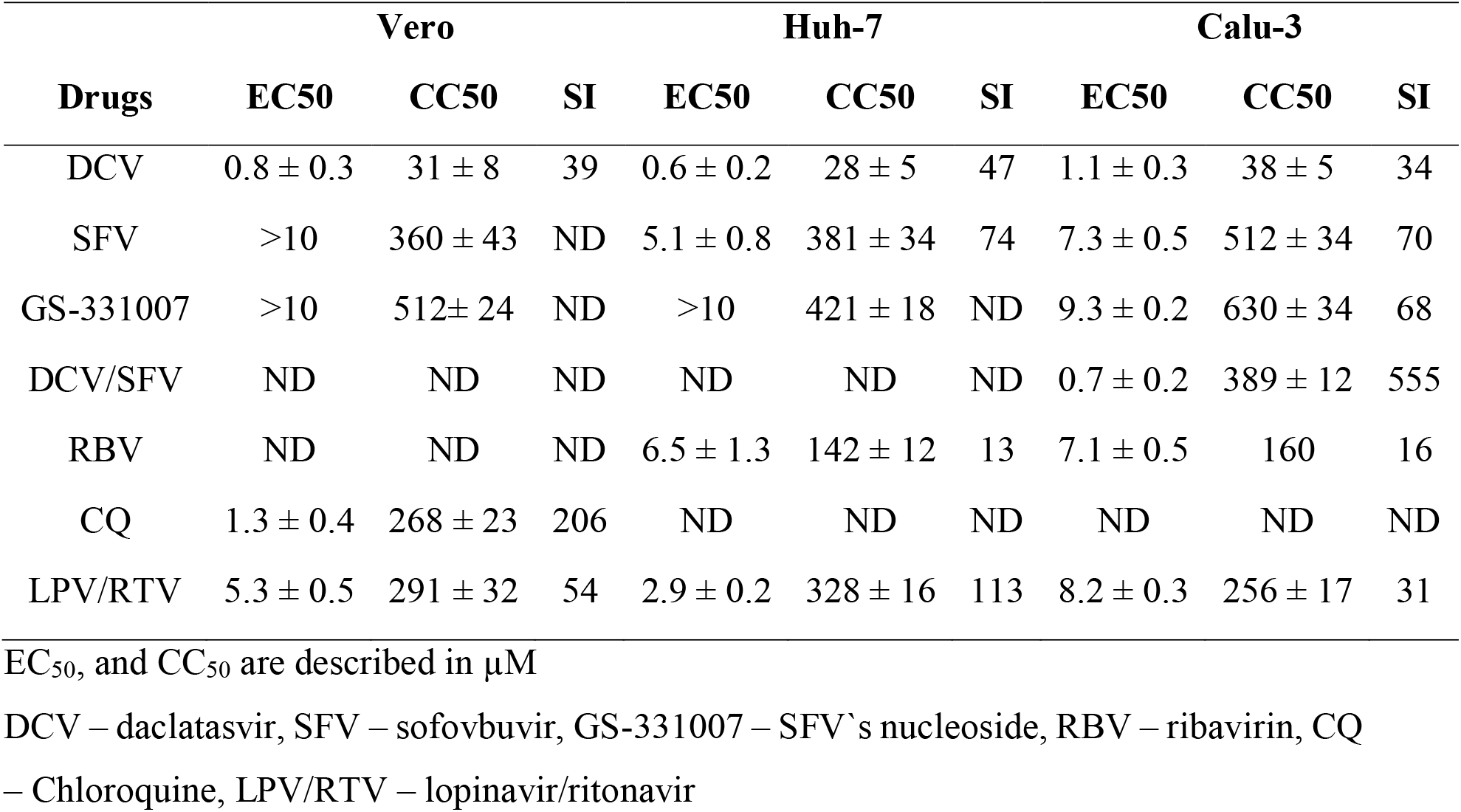
The pharmacological parameters of SARS-CoV-2 infected cell in the presence of DCV and SFV.

SARS-CoV-2 susceptibility to SFV in Huh-7 and Calu-3 cells was lower compared to DCV(Figure 1B and D and Table 1). Because vero cells poorly activate SFV to its active triphosphate, SFV did not affect SARS-CoV-2 replication in these cells. Similarly, to what was observed for DCV, quantification of SFV’s antiviral activity by PFUs was more sensitive than by viral RNA quantification in the supernatant fraction (Figure S2). DCV was at least 7-times more potent than SFV in HuH-7 and Calu-3 cells (Table 1).

SFV’s nucleoside metabolite (GS-331007) was also tested for anti-SARS-CoV-2 activity. GS-331007 was inactive in Vero cells and less active than SFV in Huh-7 cells (Figure 1 and Table 1). Curiously, in respiratory cells, GS-331007 presented a moderate anti-SARS-CoV-2 activity, similar to that of SFV (Figure 1 and Table 1).

Given that SARS-CoV-2 replication in Calu-3 cells appeared to be more sensitive to antiviral activity, this cell line was used to assess the combination of SFV and DCV. SFV/DCV combination was used at a ratio of 1:0.15 ratio, in accordance with its dose ratio for HCV-positive patients (400 mg SFV plus 60 mg DCV). In this assessment of the interaction, the potency of SFV increased 10-fold in the presence of suboptimal DCV concentrations (Figure 1C and E and Table 1).

DCV was demonstrated to be 1.1- to 4-fold more potent than the positive controls CQ, LPV/RTV and ribavirin (RBV) (Figures 1 and Table 1), whereas SFV potency was similar to that of RBV in HuH-7 and Calu-3 cells (Figures 1 and Table 1). However, the selectivity index (SI = CC_50_/EC_50_) for SFV was 4.6-fold superior to RBV, because of SFV`s lower cytotoxicity (Table 1). None of the studied drugs were more potent than RDV (Figure 1 and Table 1).

These data demonstrate that SARS-CoV-2 is susceptible to DCV and SFV *in vitro*, with a higher potency demonstrated for DCV.

### 2.2) Protective effect of SFV and DCV in non-permissive cells

Although productive replication in neurons and monocytes was not observed (Figure S1), infection of these cells is known to be associated with neuro-COVID-19[25] and cytokine storm[26], respectively. Therefore, these cell types may be important targets for repurposed antiviral drugs.

SFV reduced SARS-CoV-2 RNA levels by 20 - 40% in NSCs, at a concentration of 1 μM (Figure 2A). Conversely, no impact of DCV on SARS-CoV-2 RNA levels were observed in NSC (Figure 2A), consistently with the other cell types assayed (Figure S2). Using the more complex system of NSC-based neurospheres, the number of tunel-positive nuclei over total nuclei as a proxy of apoptotic cells was assessed. SFV completely prevented SARS-CoV-2-induced apoptosis (Figure 2B), whereas benefits of DCV in this system were limited.

In SARS-CoV-2-infected human primary monocytes, 1 μM DCV reduced viral RNA levels/cell (Figure 3A), whereas SFV was inactive. DCV also reduced the SARS-CoV-2-induced enhancement of TNF-α and IL-6 (Figure 3B and C). These data provide further evidence for a putative benefit in COVID-19 with the investigated HCV DDAs if target concentrations can be achieved in patients.

**Figure 2.**
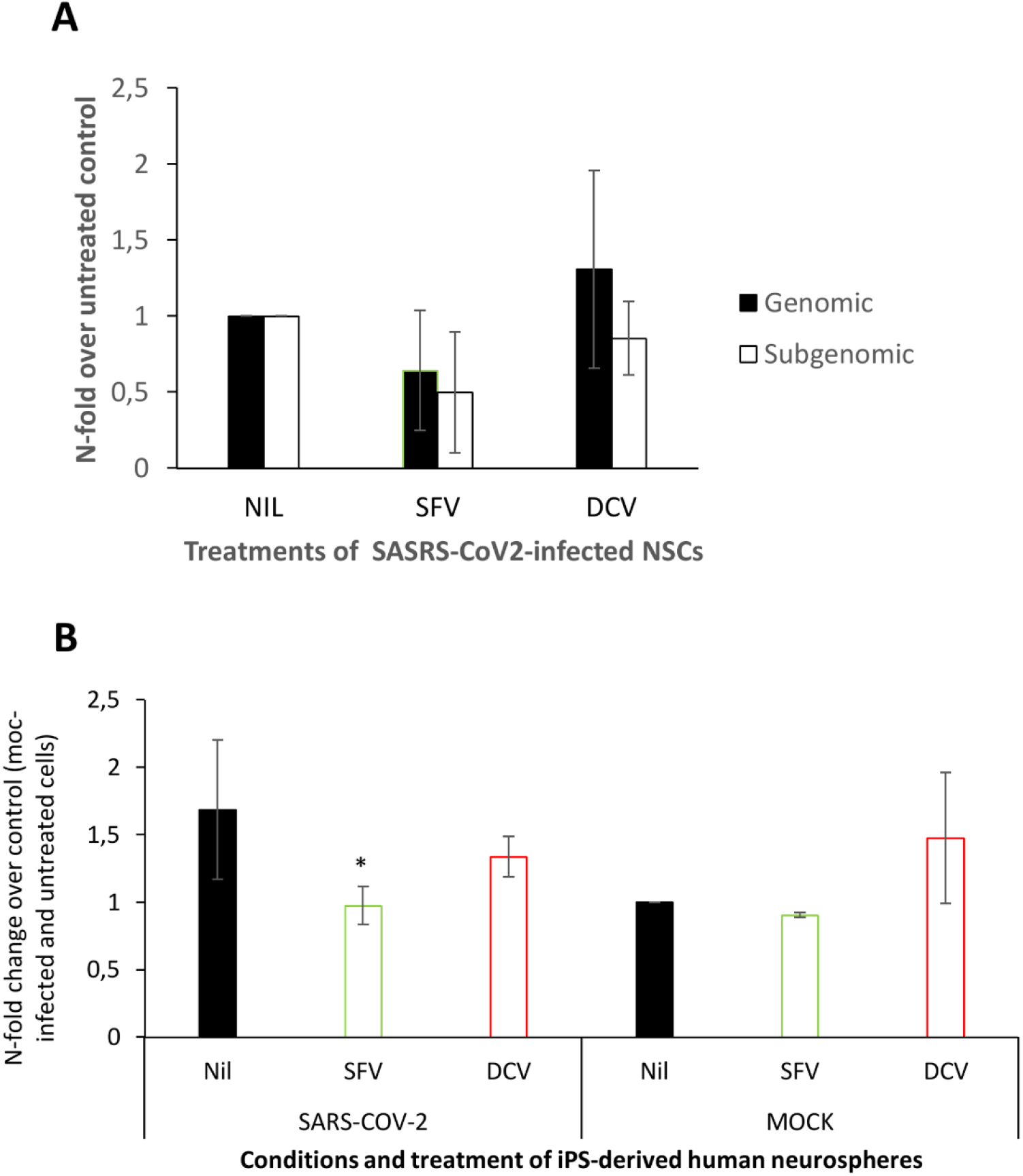
Sofosbuvir (SFV) inhibits SARS-CoV-2 replication in human iPS cell-derived NSCs. (A) NSCs were infected at MOIs of 0.1 and treated with 1 μM of SFV or daclatasvir (DCV). After 5 days, the culture supernatants were collected, and the virus was quantified by RNA levels using RT-PCR. (B) NSCs in spheroid format were labeled for Tunel and DAPI after 5 days post-infection. The data represent means ± SEM of three independent experiments. * indicates *P* < 0.05 for the comparison between the SARS-CoV-2-infected cells untreated (nil) vs treated with SFV.

**Figure 3.**
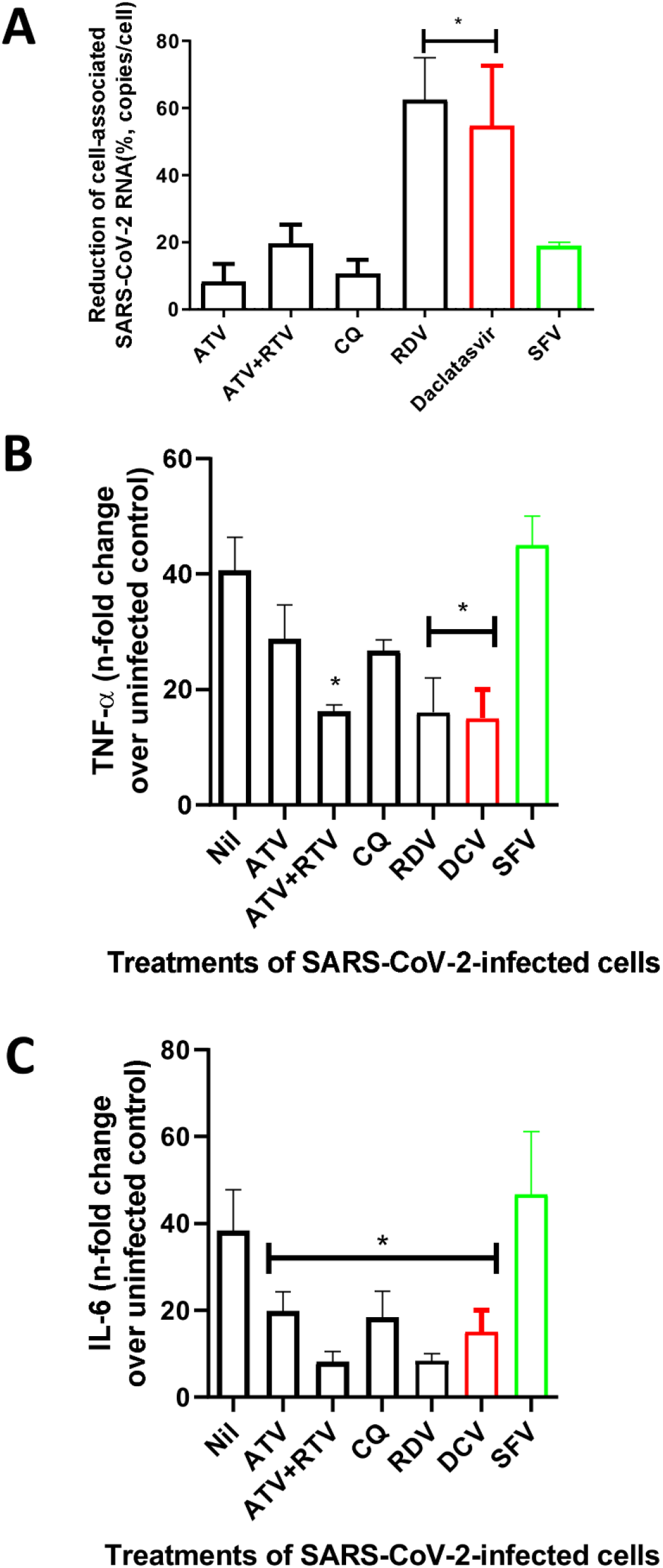
Daclatasvir (DCV) impairs SARS-CoV-2 replication and cytokine storm in human primary monocytes. Human primary monocytes were infected at the MOI of 0.01 and treated with 1 μM of daclatasvir (DCV) sofosbuvir (SFV), chloroquine (CQ), atazanavir (ATV) or atazanavir/ritonavir (ATV+RTV). After 24h, cell-associated virus RNA loads (A), as well as TNF-α (B) and IL-6 (C) levels in the culture supernatant were measured. The data represent means ± SEM of experiments with cells from at least three healthy donors. Differences with *P* < 0.05 are indicates (*), when compared to untreated cells (nil) to each specific treatment.

SFV and DCV cooperatively target virus replication in cells from different anatomical sites, preventing SARS-CoV-2-mediated neuronal cell death and the increase of pro-inflammatory mediators.

### 2.3) DCV and SFV may target different events during SARS-CoV-2 RNA synthesis

The observation that suboptimal concentrations of DCV augmented antiviral activity of SFV (Figure 1C and F) may indicate that they target different processes during viral replication. As a nucleotide analog, SFV was described to competitively inhibit the SARS-CoV-2 RNA polymerase[22]. In HCV, DCV blocks the multi-functional protein NS5A, also suggesting these agents target different mechanisms within the SARS-CoV-2 life cycle. To gain insights on the temporality of DCV`s activity against SARS-CoV-2, Vero cells were infected at MOI of 0.01 and treated at different timepoints, with DCV at 2-fold its EC_50_. This time-of-addition assay demonstrated that DCV treatment could be efficiently postponed up to 4 h, similarly to RBV, a pan-RNA polymerase inhibitor (Figure 4A). These results suggest that inhibition of viral RNA synthesis is the limiting event targeted by DCV.

**Figure 4.**
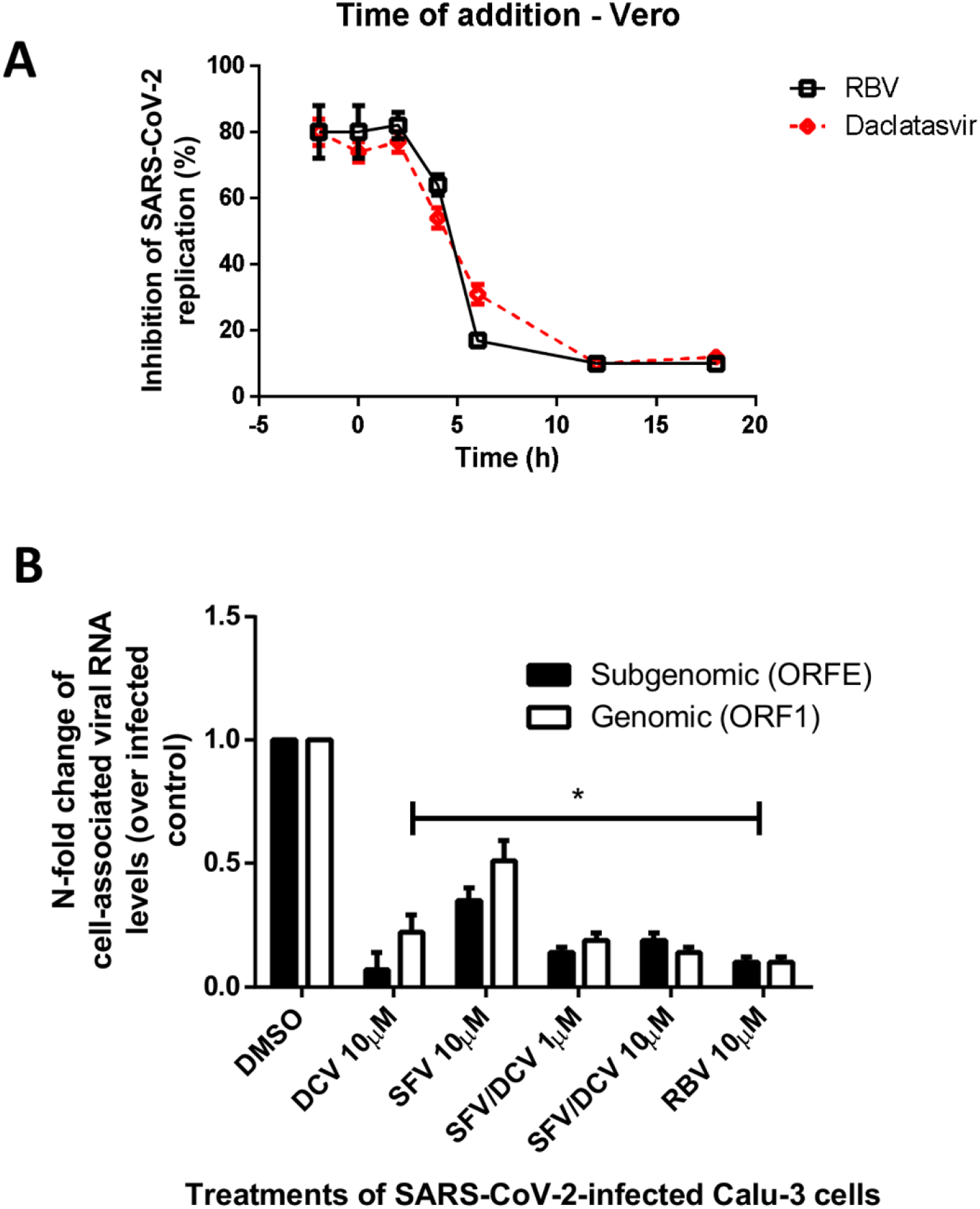
Daclatasvir (DCV) and sofosbuvir (SFV) reduced SARS-CoV-2 associated RNA synthesis. (A) To initially understand the temporal pattern of inhibition promoted daclatasvir, we performed by Time-of-addition assays. Vero cells were infected with MOI 0f 0.01 of SARS-CoV-2 and treated with daclatasvir or ribavirin (RBV) with two-times their EC_50_ values at different times after infection, as indicated. After 24h post infection, culture supernatant was harvested and SARS-CoV-2 replication measured by plaque assay. (B) Next, Calu-3 cells (5 × 10^5^ cells/well in 48-well plates), were infected with SARS-CoV-2 at MOI of 0.1, for 1h at 37 °C. Inoculum was removed, cells were washed and incubated with fresh DMEM containing 2% fetal bovine serum (FBS) and the indicated concentrations of the daclatasvir, SFV or ribavirin (RBV) at 10 μM. After 48h, cells monolayers were lysed, total RNA extracted and quantitative RT-PCR performed for detection of ORF1 and ORFE mRNA. The data represent means ± SEM of three independent experiments. * P< 0.05 for comparisons with vehicle (DMSO). # P< 0.05 for differences in genomic and sub-genomic RNA.

To confirm the rational that both SFV and DCV inhibit viral RNA synthesis in physiologically relevant cells, intracellular levels of SARS-CoV-2 genomic and subgenomic RNA were measured in type II pneumocytes, Calu-3 cells. A two-fold higher inhibition of viral RNA synthesis was observed for DCV compared to SFV (Figure 4B), when both were tested at 10 μM. SFV/DCV cooperatively inhibited SARS-CoV-2 RNA synthesis, even at 1μM, also supporting different targets for each agent during replicase activity.

Molecular docking methods were applied to predict the complexes with lowest energy interactions between the SARS-CoV-2 RNA polymerase and the active metabolite of SFV as well as DCV. The SFV active metabolite and DCV presented rerank score values of −74.09 a.u. and −84.64 a.u., respectively. In addition, the hydrogen bonds (H-bonds), attractive electrostatic, and steric interactions were mapped using a ligand-map algorithm[27]. The SFV active metabolite was predict to interact via hydrogen bonds (H-bond) with Arg553, Cys622, Asp623, and Asn691 residues and with U20 RNA nucleotide (H-bond interaction energy = −3.50 u.a.), also presenting electrostatic interactions with Lys551, Arg553, and with the two Mg^2+^ ions (electrostatic interaction energy = −13.14 u.a.), as described by Gao coworkers[21], and steric interactions with Arg553, Cys622, Asp623, and Asn691 residues (steric interaction energy = −74.09 u.a.) (Figure 5A and C). Furthermore, these predictions indicated that DCV may interact with viral RNA in the cleft of the SARS-CoV-2 RNA polymerase (Figure 5B and D), with anchoring through H-bonds with Tyr546 and Thr687 residues, and with U9 RNA nucleotide (H-bond interaction energy = 3.68 u.a.), and also showing steric interactions with Tyr546 and Thr687 residues (steric interaction energy = −84.64 u.a.) (Figure 5B and D).

**Figure 5.**
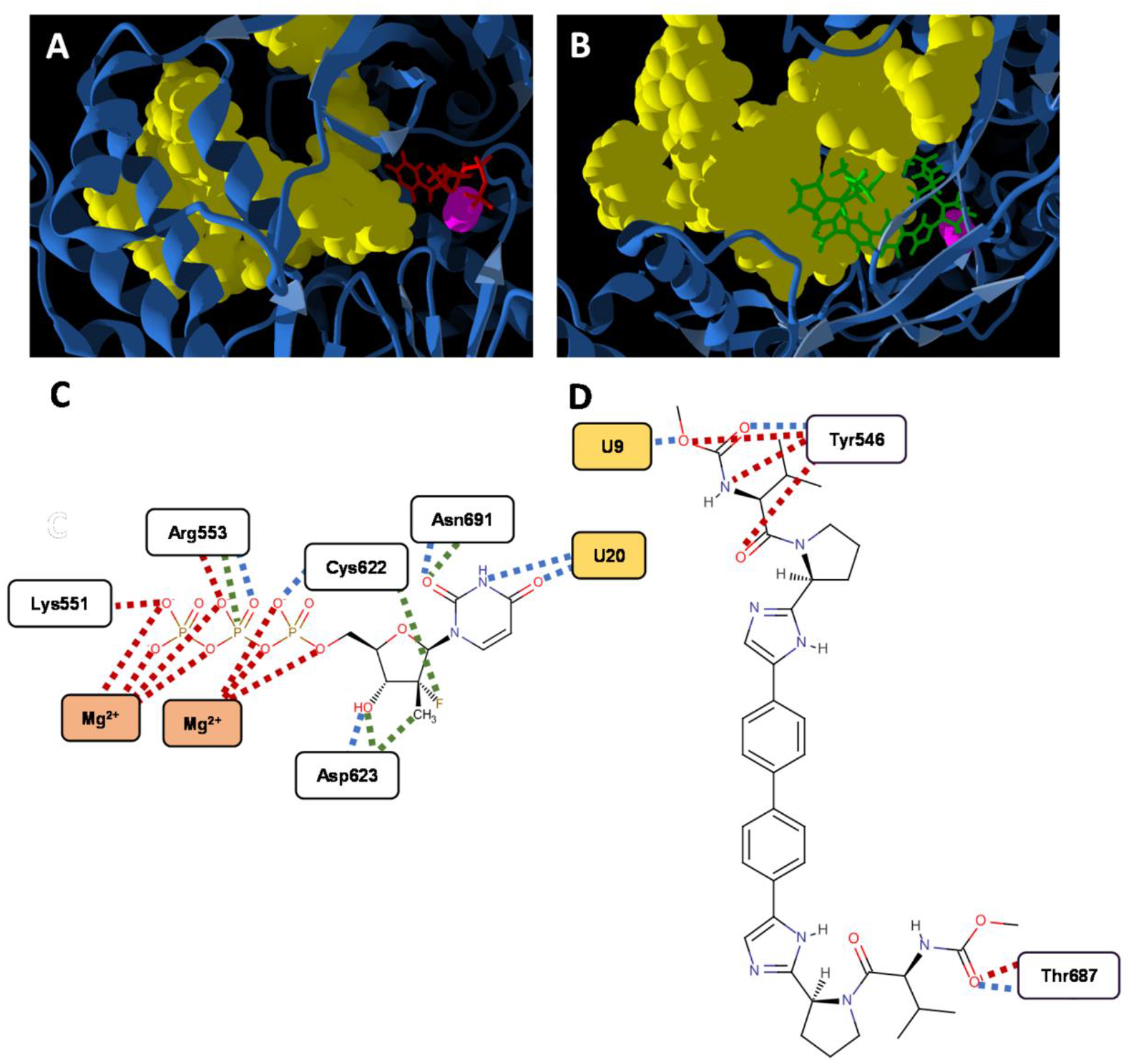
(**A**) Cartoon representation of SARS-Cov-2 RNA polymerase (nsp12; blue) with RNA template (yellow), and Mg^2+^ (pink) ions, in CPK representation, complexed to the active metabolite of sofosbuvir (SFV; red) (**A**) and daclatasvir (**B**). Schematic representations of the hydrogen bonds (H-bonds; blue dashed lines), attractive electrostatic interactions (red dashed lines), and steric interactions (green dashed lines) present in the nsp12-SFV (**C**) and nsp12-daclatasvir (**D**) complexes. The nsp12 residues, RNA nucleotides, and Mg^2+^ ions are represented by white, yellow, and orange rectangles.

### 2.4) DCV effect on SARS-CoV-2 RNA

Predictions from molecular modeling and data from in vitro phenotypic assays suggested that DCV could target SARS-CoV-2 RNA synthesis. Therefore, a melting curve of extracted viral RNA was generated to assess whether DCV could affect the virus RNA folding. SARS-CoV-2 RNA displays secondary structures throughout its sequence, which are important during viral replication and trascription[28], which can be monitored through melting curve analysis using a regular real time thermocycler. The thermal melting profiles of the RNA and RNA/DCV complexes, obtained by varying the temperature, showed concentration-dependent effects favoring denaturation of the nucleic acid at low temperatures (Figure 6A and B).

**Figure 6.**
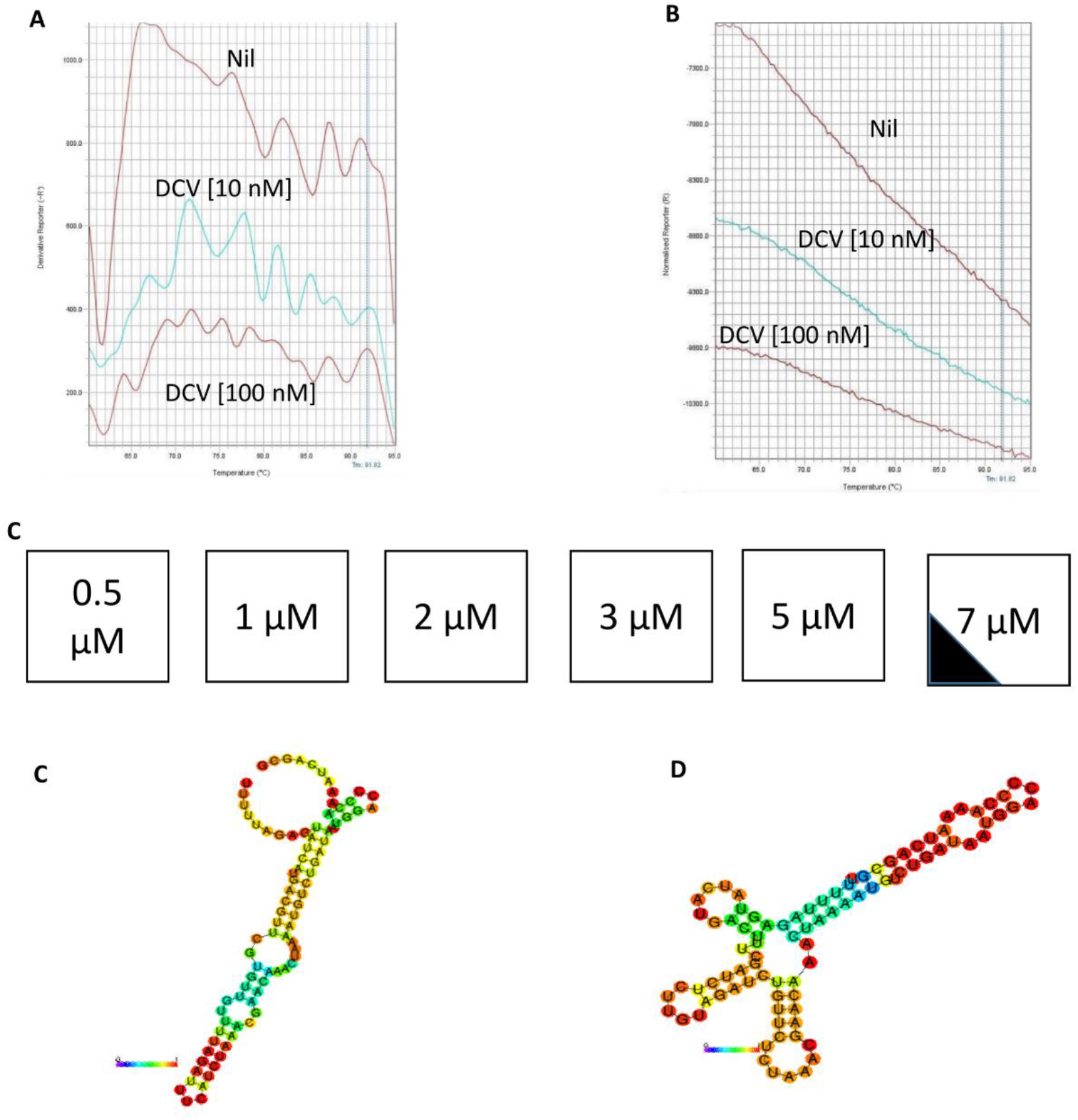
Daclatasvir (DCV) favors SARS-CoV-2 RNA unfold. A total of 10 ng of SARS-CoV-2 RNA was incubated with 10 or 100 nM of DCV during a standard melting curve in the presence of picogreen, derivative (A) and normalized (B) reports are presented. (C) the scheme represent the percentage of wild-type (WT; white) and mutant (black) virus after growing SARS-CoV-2 in Vero Cells at a MOI 10 times higher than used in other experiments, 0.1, and sequentially treated with sub-optimal doses of DCV. Each passage was done after 2-4 days pos-infection, when cytopathic effect was evident. Virus RNA was unbiased sequenced using a MGI-2000 and a metatrasncriptomic approach was employed during the analysis. WT (D) and mutant (E) SARS-CoV-2 secondary RNA structure encompassing the nucleotides 28169-28259 are presented.

In order to investigate further, it was hypothesized that continuous culture of the virus in the presence do DCV may result in mutations in the SARS-CoV-2 RNA which change the pattern of secondary structure. Following two months successive passage of the virus in Vero cells at the MOI of 0.1 in the presence of increasing concentrations, a 30% mutant subpopulation was detected in the presence of 7 μM DCV (Figure 6C). A putative secondary structure at positions 28169-28259 of the SARS-CoV-2 genome was changed in the mutant virus (yielded in the presence of DCV) in comparison to wild-type (SARS-CoV-2 virus grown in parallel without treatment) (Table 2, Figure 6D and E, genbank #MT827075, MT827190, MT827872, MT827940, MT827074, MT827202, MT835026, MT835027, MT835383, SRR12385359 and its coverage in Figure S3). The positions 28169-28259 are located at the junction between ORF8 and N gene; thus, the change in the shape of the secondary RNA structure may prevent the binding of specific components required for the transcription of these genes (Figure 6D and E). Moreover, the low sequence identity of the mutant with SARS-CoV-2 genomes in genbank suggests that it may be unlikely that mutant virus possesses adequate fitness (Table 2), which is in line with the observed reduction in virus infectious titers.

**Table 2.**
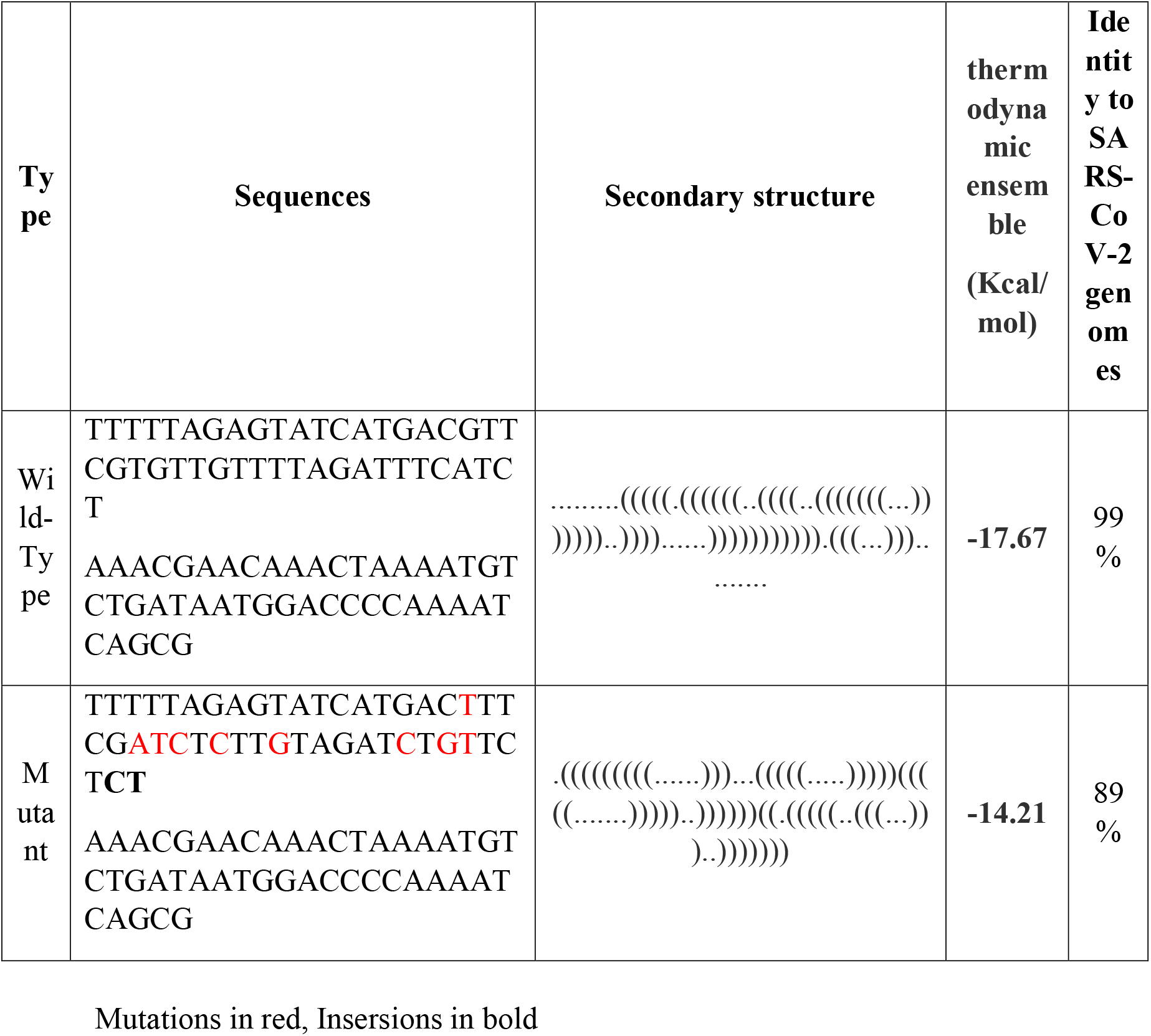
Genetic and biochemical characteristics of the DCV-mutant SARS-CoV-2.

### 2.5) Physiologically based pharmacokinetic (PBPK) modelling for DCV

A recent analysis of drugs proposed for repurposing as SARS-CoV-2 antiviral medicines revealed that very few of the proposed candidates achieved their target concentrations after administration of approved doses to humans [29]. Moreover, there have been several recent calls to integrate understanding of pharmacokinetic principles into COVID-19 drug prioritization[30–32]. Initial assessment of the plasma pharmacokinetics of SFV indicated that the concentrations able to inhibit SARS-CoV-2 replication in vitro were unlikely to be achievable after approved doses. However, inhibitory DCV concentrations were close to those achieved following administration of its approved HCV dose. Therefore, Physiologically based pharmacokinetic (PBPK) modelling was used to estimate the dose and schedule of this drug to maximize the probability of success for COVID-19.

PBPK model validation against various single and multiple oral doses of DCV had a ratio <2 between mean simulated and observed values and a summary of this shown in supplementary tables S1 and 2. The average absolute fold error (AAFE) values for the observed vs simulated plasma concentration – time curve for a single 100 mg dose and multiple 60 mg OD doses were 0.92 and 0.76, respectively, and are shown in supplementary figure S4 and S5. Thus, the known pharmacokinetic values and plots are in the agreeable range for the DCV PBPK model to assumed as validated.

Supplementary figures S6 and S7 show the C_24_ values for various BID and TID dose simulations, and 540 mg BID and 330 mg TID were shown to satisfy systemic concentrations above the EC_90_ for at least 90% of the simulated population. Optimal dose was identified to be 330 mg TID as this dosing regimen requires lower dose per day than 540 mg BID. A comparison between 60 mg TID and 330 mg TID daclatasvir is shown in Figure 7 that satisfy C_24_ for EC_50_ (0.8 μM, 591 ng/ml) and EC_90_ respectively for treatment of SARS-CoV-2.

**Figure 7.**
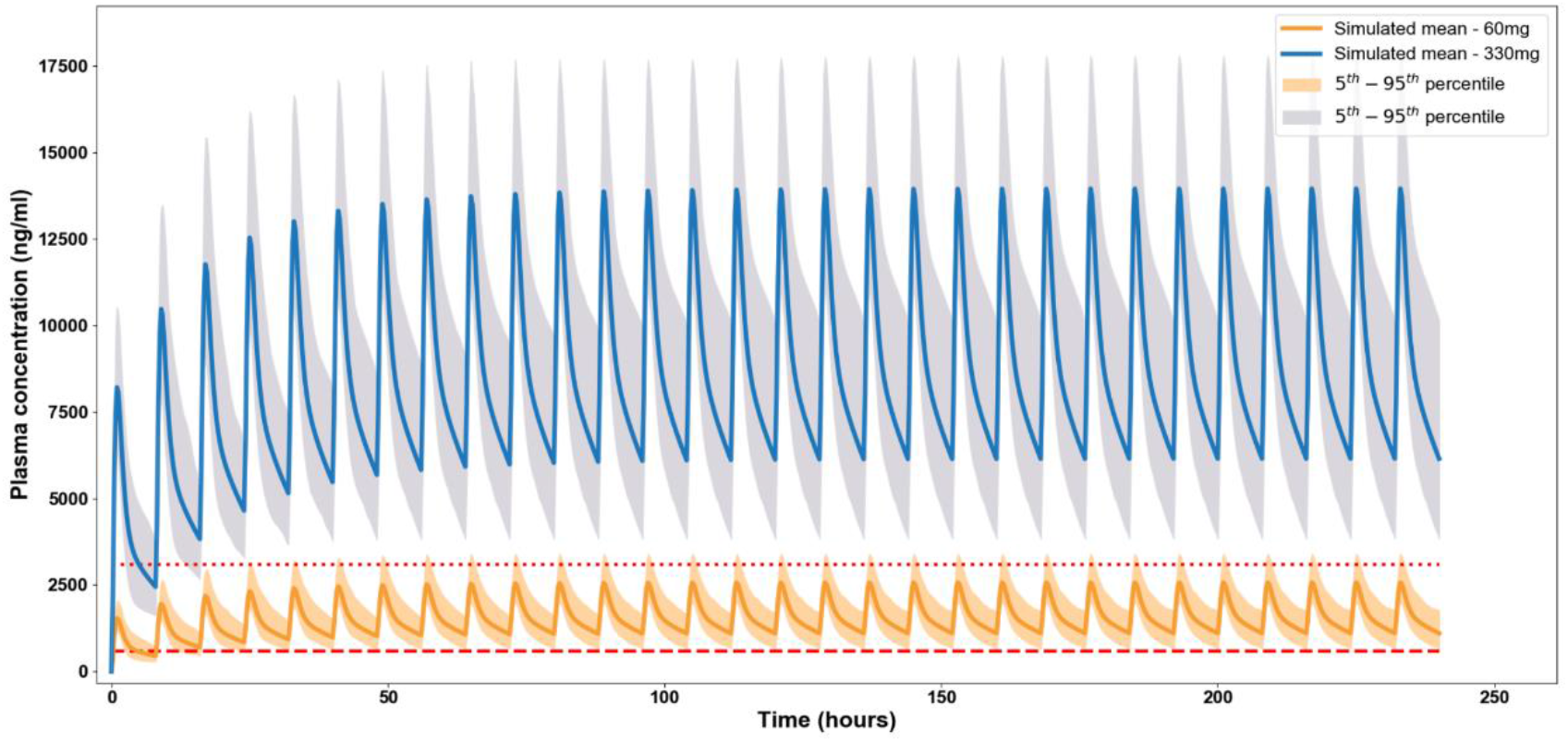
Predicted daclatasvir plasma concentration for multiple 60 mg and 330 mg TID doses. The dotted and the dashed lines represent the EC_90_ and EC_50_ values of daclatasvir for SARS-CoV-2.

## 3) Discussion

The COVID-19 pandemic continues to present a major concern to global health, and is the most significant economic threat in decades[33]. Less than 8 months after the outbreak in Wuhan, China, the WHO recorded more 750,000 deaths worldwide^1^. SARS-CoV-2 is the third highly pathogenic coronavirus that emerged in the two decades of the 21^st^ century, following SARS-CoV and MERS-CoV[1]. SARS-CoV-2 actively replicates in type II pneumocytes, leading to cytokine storm and the exacerbation of thrombotic pathways[26,34,35]. Besides the virus-triggered pneumonia and sepsis-like disease associated with severe COVID-19, SARS-CoV-2 may reach the central nervous system[25] and liver[36]. Early blockage of the natural clinical evolution of infection by antivirals will likely prevent the disease progression to severe COVID-19[26,34,35]. Indeed, clinical studies providing early antiviral intervention accelerated the decline of viral loads and slowed disease progression[7,8]. The decrease of viral loads is likely to be a critical laboratory parameter, because lowering viral shedding may protect the individual and reduced transmissibility is likely to have population-level benefits.

To rapidly respond to unfolding pandemics, the cataloguing of preclinical data on susceptibility of SARS-CoV-2 to approved drugs is of paramount importance, and provides opportunities for rational selection of promising products for evaluation in clinical trials [37]. The investigators used this approach during ZIKV, YFV, and CHIKV outbreak in Brazil, and demonstrated susceptibility of these viruses to SFV [13–16,38]. SFV and DCV are considered safe and well tolerated anti-HCV therapies that are orally bioavailable. The presented work demonstrates: i) SARS-CoV-2 is susceptible to DCV, ii) DCV/SFV co-treatment show cooperative antiviral effect on SARS-CoV-2 replication in respiratory cells; iii) SFV and DCV prevented virus-induced neuronal apoptosis and release of cytokine storm-related mediators in monocytes, respectively; iv) DCV and SFV inhibited independent events during RNA synthesis; v) DCV favors the unfold of SARS-CoV-2 secondary RNA structures, and vi) target concentration of DCV set by the *in vitro* activities are within the range that may be achievable in humans.

In the 9.6 kb genome of HCV, the gene *ns5a* encodes for a multifunctional protein. The protein NS5A possesses motifs involved with lipid, zinc and RNA binding, phosphorylation and interaction with cell signaling events[10]. In other viruses, with less compact genomes, the functions and motifs present in NS5A are distributed to other proteins. For instance, in SARS-CoV-2, its 29 kb genome encodes for nsp3, with zinc motif; nsp4 and 5, with lipidic binding activity; nsp7, 8, 12, 13 and 14 able to bind RNA[11]. Although there is not a specific orthologue of NS5A in the SARS-CoV-2 genome, their activities may be exerted by multiple other proteins. DCV inhibited the production of infectious SARS-CoV-2 titers with EC_50_ values ranging from 0.6 to 1.1 μM across different cell types, including pneumocytes. Curiously, DCV`s antiviral activity was not exhibited when virus replication was accessed by quantifying viral RNA loads. Our sub-sequential analysis illustrated that DCV mechanism of action could be, at least in part, associated with targeting viral RNA secondary structures, in line with the observation of lower infectivity in the absence of viral RNA decline in culture supernatant. SARS-CoV-2 possesses RNA pseudoknots that could contribute to the transcription processes[28], and DCV-associated denaturation of these structures could limit viral RNA polymerase activity. This already impaired catalysis may promote cooperative activity of SFV.

With relevance to SFV, the homology of the new-2019-CoV and HCV orthologue enzyme were confirmed [21]. In enzyme kinetic assays with SARS-CoV-2 nsp7, 8 and 12, the SARS-CoV-2 RNA polymerase complex, SFV-triphosphate, the active metabolite, competitively acts as a chain terminator[22,23]. Similarly, RBV-, favipiravir-and RDV-triphosphate also target SARS-CoV-2 RNA elongation[22,23]. Indeed, SFV reduced the RNA synthesis in SARS-CoV-2-infected cells able to convert the pro-drug to its active triphosphate, such as hepatoma cells. This activation process requires a multi-stage pathway in which hydrophobic protections in the SFV monophosphate are removed by the cellular enzymes CatA, CES1 and HINT, with subsequent followed by engagement of nucleoside monophosphate and diphosphate kinase [17]. According to the Human Protein Atlas, these enzymatic entities are also found in the respiratory tract[18–20]. Indeed, we found that SARS-CoV-2 replication could be inhibited by SFV at high concentration, not only in hepatoma cells – but also in Calu-3 type II pneumocytes. Interestingly, RDV, which shares structural characteristics with SFV, such as to be converted from the ProTide/prodrug to active metabolite, is active in the respiratory tract[39]. Moreover, there is a body of evidence suggesting that the ProTide phospharamidate protections would be dispensable from RDV in respiratory cells because the nucleoside analog, GS-441524, is active against human and feline CoV [39–41]. Since there are open questions on the efficiency in which respiratory cells convert nucleosides to nucleotides, the nucleoside version of SFV (GS-331007) was tested against SARS-CoV-2. GS-331007 was virtually inactive in all cell, types except for Calu-3, in which it exerted similar activity to SFV. Importantly, GS-331007 has broader distribution in anatomical compartments than SFV, which may be important in the context of anatomical target-site activity.

Considering that DCV could favor RNA denaturation, conformational changes in the viral RNA template/primer dimer at nsp12 active site may limit efficiency or processing by this enzyme. Since SARS-CoV-2 RNA polymerase kinetics is impaired by DCV, SFV could be less impacted by hindrance via amino acid Asp623[22] in this enzyme. This hypothesis warrants further investigation to confirm the mechanistic-basis for the possible cooperation between SFV and DCV *in vitro* model, and clinically if observations from recent trials are confirmed[42].

SFV was able to prevent apoptosis in human neurons, whereas DCV prevented the enhancement of IL-6 and TNF-α levels in human monocytes. These secondary mechanisms may also support cooperativity between SFV and DCV, because neurological SARS-CoV-2 infection and cytokine storm are associated with poor clinical outcomes[25,26]. Another study also reported that SFV could be protective against neuro-COVID *in vitro*[24]. However, the authors analyzed only a single dose of 20 μM, which greatly exceeds the concentrations achieved by SFV after approved dosing to humans [17]. Here, neuroprotection is demonstrated to be promoted by SFV at 1 μM, which is closer to physiological concentrations [17].

Based upon targets set by the *in vitro* pharmacological activity of DCV, PBPK modelling indicated that systemic concentrations able to inhibit SARS-CoV-2 may be achievable in humans. Dose escalation may be needed to provide fully suppressive concentrations across the entire dosing interval, as has been shown to be needed for other viruses. However, the validity of such an approach would require careful assessment of safety and tolerability through phase I evaluation of the higher doses. Furthermore, the prerequisite pharmacokinetic-pharmacodynamic relationships for successful anti-SARS-CoV-2 activity are yet to be unraveled, and will likely require better understanding of the target-site penetration and free drug concentrations in matrices that recapitulate relevant compartments. Notwithstanding, the approved dose of DCV (60mg OD) is low in relationship to other antiviral agents, and the PBPK model provides posologies that may be reachable in dose-escalation trials.

In summary, effective early antiviral interventions are urgently required for the SARS-CoV-2 pandemic to improve patient clinical outcomes and disrupt transmission at population level. The presented data for two widely available anti-HCV drugs, particularly for DCV, provide a rational basis for further validation of these molecules for anti-SARS-CoV-2 interventions.

## 4) Material and Methods

### 4.1. Reagents

The antivirals RDV and LPV/RTV (4:1 proportion) was pruchased from Selleckhem (https://www.selleckchem.com/). Chloroquine and ribavirin were received as donations from Instituto de Tecnologia de Fármacos (Farmanguinhos, Fiocruz). DCV and SFV were donated by Microbiologica Química-Farmacêutica LTDA (Rio de Janeiro, Brazil). ELISA assays were purchased from R&D Bioscience. All small molecule inhibitors were dissolved in 100% dimethylsulfoxide (DMSO) and subsequently diluted at least 10^4^-fold in culture or reaction medium before each assay. The final DMSO concentrations showed no cytotoxicity. The materials for cell culture were purchased from Thermo Scientific Life Sciences (Grand Island, NY), unless otherwise mentioned.

### 4.2. Cells and Virus

African green monkey kidney (Vero, subtype E6) and, human hepatoma (Huh-7), human lung epithelial cell lines (A549 and Calu-3) cells were cultured in high glucose DMEM and human hepatoma lineage (Huh-7) in low glucose DMEM medium, both complemented with 10% fetal bovine serum (FBS; HyClone, Logan, Utah), 100 U/mL penicillin and 100 μg/mL streptomycin (Pen/Strep; ThermoFisher) at 37 °C in a humidified atmosphere with 5% CO2.

Human primary monocytes were obtained after 3 h of plastic adherence of peripheral blood mononuclear cells (PBMCs). PBMCs were isolated from healthy donors by density gradient centrifugation (Ficoll-Paque, GE Healthcare). PBMCs (2.0 × 10^6^ cells) were plated onto 48-well plates (NalgeNunc) in RPMI-1640 without serum for 2 to 4 h. Non-adherent cells were removed and the remaining monocytes were maintained in DMEM with 5% human serum (HS; Millipore) and penicillin/streptomycin. The purity of human monocytes was above 95%, as determined by flow cytometric analysis (FACScan; Becton Dickinson) using anti-CD3 (BD Biosciences) and anti-CD16 (Southern Biotech) monoclonal antibodies.

NSCs derived from human iPS cells were prepared as previously described[43]. N3D human neurospheres were generated from 3 × 10 6 NSCs/well in a 6-well plate orbital shaking at 90 rpm and were grown in NEM supplemented with 1×N2 and 1×B27 supplements. After 7 days in culture, neurospheres or NSC were infected at MOI 0.1 for 2h at 37 °C. NSCs were washed, neurospheres inoculum were aspirate, and fresh medium containing the compounds was added. Neural cells were observed daily for 5 days after infection. Cell death was measured by tunel approach and virus levels in the supernatant quantified by RT-PCR.

SARS-CoV-2 was prepared in Vero E6 cells at MOI of 0.01. Originally, the isolate was obtained from a nasopharyngeal swab from a confirmed case in Rio de Janeiro, Brazil (GenBank #MT710714; Institutional Review Broad approval, 30650420.4.1001.0008). All procedures related to virus culture were handled in a biosafety level 3 (BSL3) multiuser facility according to WHO guidelines. Virus titers were determined as plaque forming units (PFU)/mL. Virus stocks were kept in −80 °C ultralow freezers.

### 4.3. Cytotoxicity assay

Monolayers of 1.5 × 10^4^ cells in 96-well plates were treated for 3 days with various concentrations (semi-log dilutions from 1000 to 10 μM) of the antiviral drugs. Then, 5 mg/ml 2,3-bis-(2-methoxy-4-nitro-5-sulfophenyl)-2*H*-tetrazolium-5-carboxanilide (XTT) in DMEM was added to the cells in the presence of 0.01% of N-methyl dibenzopyrazine methyl sulfate (PMS). After incubating for 4 h at 37 °C, the plates were measured in a spectrophotometer at 492 nm and 620 nm. The 50% cytotoxic concentration (CC_50_) was calculated by a non-linear regression analysis of the dose–response curves.

### 4.4. Yield-reduction assay

Unless otherwise mentioned, Vero E6 cells were infected with a multiplicity of infection (MOI) of 0.01. HuH-7, A549 and Calu-3 were infected at MOI of 0.1. Cells were infected at densities of 5 × 105 cells/well in 48-well plates for 1h at 37 °C. The NSCs (20 × 10³ cells/well in a 96-well plate) were infected at MOI of 0.1 for 2 h at 37 °C. The cells were washed, and various concentrations of compounds were added to DMEM with 2% FBS. After 24 (Vero E6), or 48h (HuH, −7, A549 and Calu-3) or 5 days (NSCs) supernatants were collected and harvested virus was quantified by PFU/mL or real time RT-PCR. A variable slope non-linear regression analysis of the dose-response curves was performed to calculate the concentration at which each drug inhibited the virus production by 50% (EC_50_).

For time-of-addition assays, 5 x 10^5^ Vero E6 cells/well in 48-well plates were infected with MOI of 0.01 for 1h at 37 °C. Treatments started from 2h before to 18h after infection with two-times EC_50_ concentration. On the next day, culture supernatants were collected and tittered by PFU/mL.

### 4.5. Virus titration

Monolayers of Vero E6 cells (2 x 10^4^ cell/well) in 96-well plates were infected with serial dilutions of supernatants containing SARS-CoV-2 for 1h at 37°C. Fresh semi-solid medium containing 2.4 % of carboxymethylcellulose (CMC) was added and culture was maintained for 72 h at 37 °C. Cells were fixed with 10 % Formaline for 2 h at room temperature and then, stained with crystal violet (0.4 %). Plaque numbers were scored in at least 3 replicates per dilution by independent readers. The reader was blind with respect to source of the supernatant. The virus titers were determined by plaque-forming units (PFU) per milliliter.

### 4.6. Molecular detection of virus RNA levels

The total viral RNA from a culture supernatants and/or monolayers was extracted using QIAamp Viral RNA (Qiagen®), according to manufacturer’s instructions. Quantitative RT-PCR was performed using GoTaq® Probe qPCR and RT-qPCR Systems (Promega) in an StepOne™ Real-Time PCR System (Thermo Fisher Scientific) ABI PRISM 7500 Sequence Detection System (Applied Biosystems).Amplifications were carried out in 25 μL reaction mixtures containing 2× reaction mix buffer, 50 μM of each primer, 10 μM of probe, and 5 μL of RNA template. Primers, probes, and cycling conditions recommended by the Centers for Disease Control and Prevention (CDC) protocol were used to detect the SARS-CoV-2[44]. The standard curve method was employed for virus quantification. For reference to the cell amounts used, the housekeeping gene RNAse P was amplified. The Ct values for this target were compared to those obtained to different cell amounts, 10^7^ to 10^2^, for calibration. Alternatively, genomic (ORF1) and subgenomic (ORFE) were detected, as described elsewhere[45].

### 4.7. Melting curve assay

The melting profiles were obtained incubating 10 ng of SARS-CoV-2 RNA with 10 or 100nM of DCV and Sybergreen (1x) (Thermo Fisher Scientific) in an StepOne™ Real-Time PCR System (Thermo Fisher Scientific) programed with default melting curve. RNA A260/280 ratio was above 1.8, consistent with consistent with high quality material.

### 4.8. Generation of mutant virus

Vero E6 cells were infected with SARS-CoV-2 at a MOI 0.1 (10-fold higher than used in the pharmacological assays) for 1h at 37 °C and then treated with sub-optimal dose of DCV. Cells were accompanied daily up to the observation of cytophatic effects (CPE). Virus was recovered from the culture supernatant, titered and used in a next round of infection in the presence of higher drug concentration. Concentrations of DCV ranged from 0.5 to 7 μM. As a control, SARS-CoV-2 was also passaged in the absence of treatments to monitor genetic drifts associated with culture. Virus RNA virus was extracted by Qiamp viral RNA (Qiagen) and quantified using Qbit 3 Fluorometer (Thermo Fisher Scientific) according to manufacters recommendations.

The virus RNA was submitted to unbiased sequence using a MGI-2000 and a metatranscriptomics approach. To do so, at least 4.2 ng of purified total RNA of each sample was used for libraries construction using the MGIEasy RNA Library Prep Set (MGI, Shenzhen, China). All libraries were constructed through RNA‐ fragmentation (250 bp), followed by reverse‐ transcription and second‐ strand synthesis. After purification with MGIEasy DNA Clean Beads (MGI, Shenzhen, China), were submitted to end‐ repair, adaptor‐ ligation, and PCR amplification steps. After purification as previously described, samples were quantified with Qubit 1X dsDNA HS Assay Kit using an Invitrogen Qubit 4.0 Fluorometer (Thermo Fisher Scientific, Foster City, CA) and homogeneously pooled (1 pmol/ pool of PCR products) and submitted to denaturation and circularization steps to be transformed into a single‐ stranded circular DNA library. Purified libraries were quantified with Qubit ssDNA Assay Kit using Invitrogen Qubit 4.0 Fluorometer (Thermo Fisher Scientific, Foster City, CA) and DNA nanoballs were generated by rolling circle amplification of a pool (40 fmol/ reaction), then quantified as described for the libraries and loaded onto the flow cell and sequenced with PE100 (100-bp paired-end reads).

Sequencing data were initially analyses in the usegalaxy.org platform. Next, aligned therough clustalW, usina the Mega 7.0 software.

### 4.9. TUNEL (Terminal deoxynucleotidyl transferase-mediated biotinylated UTP Nick End Labelling)

Nuclei from human neurospheres were obtained by isotropic fractionation and plated in 384 plates coated with 0.1 mg/ml poly-L-lysine. Cell death was detected by Apoptag® Red in situ apoptosis detection kit (Merck, catalog # S7165) which labels apoptotic cells, based on staining of 3’-OH termini of DNA strand breaks with rhodamine (red fluorescence), staining was performed according to manufacturer’s instructions. Nuclei were labelled with 0.5 μg/mL 40-6-diamino-2-phenylindole (DAPI) for 10 minutes. Nuclei were washed with PBS, mounted with glycerol and analyzed in an Operetta high-content imaging system with a 40x objective and high numerical apertures (NA) (PerkinElmer, USA). The data was analyzed using the high-content image analysis software Harmony 5.1 (PerkinElmer, USA). Twelve independent fields were evaluated from duplicate wells per experimental condition.

### 4.10. Molecular docking

The structures of the active metabolite of SFV and daclatasvir were constructed and optimized by the semi-empirical method RM1, using the Spartan’10 software. The crystal structure of the SARS-Cov-2 nsp12 (PDB code: 7BV2) was extracted from the Protein Data Bank[21].

The molecular docking procedure was performed using the Molegro Virtual Docker 6.0 software (MVD) [27], which uses a heuristic search algorithm that combines differential evolution with a cavity prediction algorithm. Thus, the MolDock Optimizer search algorithm was used with a minimum of 30 runs, the largest enzyme cavity (1446.4 Å^3^) was chosen as the center of the search space, and the parameter settings were: population size = 100; maximum iteration = 2000; scaling factor = 0.50; offspring scheme = Scheme 1; termination scheme = variance-based; and crossover rate = 0.90. The complexes of the lowest energy were selected using the rerank scoring function and, then, analyzed also using MVD.

### 4.11 PBPK model

DCV whole-body PBPK model was constructed in Python 3.5 (in PyCharm 20.1.2 (Communtiy edition) using packages – numpy v1.18.5, scipy v1.0.1 and matplotlib v2.1.2) which consists of various compartments representing all the organs and tissues of the body. The drug physicochemical parameters for daclatasvir were presented in supplementary table S1 obtained from various literature sources. The PBPK model was constructed based on few assumptions: 1) uniform and instant distribution across a given tissue, 2) no reabsorption from the colon and 3) the model was blood-flow limited. The simulated data in humans is computer generated, therefore no ethical approval was required for this study.

### 4.12. Model development

The model was simulated using a population of one hundred virtual healthy individuals (50% female) between 20-60 years and having weight and height as provided by the US national health statistics reports[46]. Organ weights and volumes, blood flow rates were obtained using anthropometric equations from literature[47,48] and the characteristics such as weight and height from US statistics[46]. A seven compartmental absorption and transit model representing the various parts of the duodenum, jejunum and ileum to capture effective absorption kinetics was used in the model. The drug was assumed to have entire administered dose in solution for absorption and completely depend on the rate kinetics involved during this process. Effective permeability of daclatasvir was scaled from apparent permeability from PAMPA (due to lack of available data, it was assumed the same in Caco-2 cells) using the *in vitro – in vivo* extrapolation[49,50] to compute the absorption rate from the small intestine.

The volume of distribution was computed using the tissue to plasma ratios computed from Rogers & Rowland [51] and a tissue to plasma partition factor (Kp factor) of 0.025 was used to adjust the volume of distribution to the literature value of 47 L[52]. A population of 100 individuals was simulated by varying the mean values with available standard deviation for each of the parameters in the model such that every simulation represents a unique individual.

### 4.13. Model validation

DCV PBPK model was validated in healthy individuals using available data in humans for various single doses – 1, 10, 25, 50, 100 and 200 mg and for various multiple doses – 1, 10, 30 and 60 mg at fasted state. Clinical data was digitised using Web Plot Digitiser® software from available plots. The model was considered validated when: 1) closeness of the simulated points to the literature data computed using absolute average fold error (AAFE) between the simulated and observed plasma concentration – time points was less than two; and 2) the mean simulated pharmacokinetic parameters - maximum concentration (C_max_) and the area under the plasma concentration-time curve (AUC) were less than two-fold from mean observed values.

### 4.14. Model simulations

For the inhibition of SARS-CoV-2, a mean target concentration (EC_90_) of 4.12 μM or 3079 ng/ml obtained from multiple *in vitro* studies was used [53]. Optimal dosing regimen for treatment of SARS-CoV-2 was identified from various BID and TID dosing regimens such that at least 90% of the simulated population have trough plasma concentration at 24 h (C_24_) over the mean target concentration with a low overall total dose per day.

### 4.15. Ethics Statement

Experimental procedures involving human cells from healthy donors were performed with samples obtained after written informed consent and were approved by the Institutional Review Board (IRB) of the Oswaldo Cruz Foundation/Fiocruz (Rio de Janeiro, RJ, Brazil) under the number 397-07. The National Review Board approved the study protocol (CONEP 30650420.4.1001.0008), and informed consent was obtained from all participants or patients’ representatives.

### 4.16. Statistical analysis

The assays were performed blinded by one professional, codified and then read by another professional. All experiments were carried out at least three independent times, including a minimum of two technical replicates in each assay. The dose-response curves used to calculate EC_50_ and CC_50_ values were generated by variable slope plot from Prism GraphPad software 8.0. The equations to fit the best curve were generated based on R^2^ values ≥ 0.9. Student’s T-test was used to access statistically significant *P* values <0.05. The statistical analyses specific to each software program used in the bioinformatics analysis are described above.

## Supporting information

SARS-CoV-2 and SOF-DAC_Updated SI

## Acknowledgments

Thanks are due to Prof. Andrew Hill from the University of Liverpool and Dr. James Freeman from the GP2U Telehealth for simulative scientific debate. Dr. Carmen Beatriz Wagner Giacoia Gripp from Oswaldo Cruz Institute is acknowledged for assessments related to BSL3 facility. Dr. Andre Sampaio from Farmanguinhos, platform RPT11M, is acknowledged for kindly donate the Calu-3. We thank the Hemotherapy Service of the Hospital Clementino Fraga Filho (Federal University of Rio de Janeiro, Brazil) for providing buffy-coats. This work was supported by Conselho Nacional de Desenvolvimento Científico e Tecnológico (CNPq), Fundação de Amparo à Pesquisa do Estado do Rio de Janeiro (FAPERJ). This study was financed in part by the Coordenação de Aperfeiçoamento de Pessoal de Nível Superior-Brasil (CAPES) - Finance Code 001. Funding was also provided by CNPq, CAPES and FAPERJ through the National Institutes of Science and Technology Program (INCT) to Carlos Morel (INCT-IDPN). Thanks are due to Oswaldo Cruz Foundation/FIOCRUZ under the auspicious of Inova program (B3-Bovespa funding). The funding sponsors had no role in the design of the study; in the collection, analyses, or interpretation of data; in the writing of the manuscript, and in the decision to publish the results.

## Author contributions

Experimental execution and analysis – CQS, NFR, JRT, SSGD, APDDS, CSS, ACF, MM, CRRP, CSF, VCS, FBS, MAF, CSGP, GV, LRQS, LGS, LVBH, TVAF, FSCB, MMB, RKRR

Data analysis, manuscript preparation and revision – CQS, NFR, JRT, MZPG NB, FAB, AO, MZG, SKR, DCBH, PTB, TMLS

Conceptualized the experiments – CQS, NFR, JRT, TMLS

Study coordination – TMLS

Manuscript preparation and revision – DCBH, PTB, TMLS

**The authors declare no competing financial interests.**

## References

1. Cui J, Li F, Shi Z-L. Origin and evolution of pathogenic coronaviruses. Nat Rev Microbiol. 2019;17: 181–192. doi:10.1038/s41579-018-0118-9

2. Dong E, Du H, Gardner L. An interactive web-based dashboard to track COVID-19 in real time. The Lancet Infectious Diseases. 2020;0. doi:10.1016/S1473-3099(20)30120-1

3. Organization WH. WHO R&D Blueprint: informal consultation on prioritization of candidate therapeutic agents for use in novel coronavirus 2019 infection, Geneva, Switzerland, 24 January 2020. 2020 [cited 29 Mar 2020]. Available: https://apps.who.int/iris/handle/10665/330680

4. Borba MGS, Val FFA, Sampaio VS, Alexandre MAA, Melo GC, Brito M, et al. Effect of High vs Low Doses of Chloroquine Diphosphate as Adjunctive Therapy for Patients Hospitalized With Severe Acute Respiratory Syndrome Coronavirus 2 (SARS-CoV-2) Infection: A Randomized Clinical Trial. JAMA Netw Open. 2020;3: e208857–e208857. doi:10.1001/jamanetworkopen.2020.8857

5. Wang Y, Zhang D, Du G, Du R, Zhao J, Jin Y, et al. Remdesivir in adults with severe COVID-19: a randomised, double-blind, placebo-controlled, multicentre trial. The Lancet. 2020;395: 1569–1578. doi:10.1016/S0140-6736(20)31022-9

6. Cao B, Wang Y, Wen D, Liu W, Wang J, Fan G, et al. A Trial of Lopinavir-Ritonavir in Adults Hospitalized with Severe Covid-19. N Engl J Med. 2020. doi:10.1056/NEJMoa2001282

7. Goldman JD, Lye DCB, Hui DS, Marks KM, Bruno R, Montejano R, et al. Remdesivir for 5 or 10 Days in Patients with Severe Covid-19. New England Journal of Medicine. 2020;0: null. doi:10.1056/NEJMoa2015301

8. Beigel JH, Tomashek KM, Dodd LE, Mehta AK, Zingman BS, Kalil AC, et al. Remdesivir for the Treatment of Covid-19 — Preliminary Report. New England Journal of Medicine. 2020;0: null. doi:10.1056/NEJMoa2007764

9. De Clercq E, Li G. Approved Antiviral Drugs over the Past 50 Years. Clin Microbiol Rev. 2016;29: 695–747. doi:10.1128/CMR.00102-15

10. Smith MA, Regal RE, Mohammad RA. Daclatasvir: A NS5A Replication Complex Inhibitor for Hepatitis C Infection. Ann Pharmacother. 2016;50: 39–46. doi:10.1177/1060028015610342

11. Gordon DE, Jang GM, Bouhaddou M, Xu J, Obernier K, O’Meara MJ, et al. A SARS-CoV-2-Human Protein-Protein Interaction Map Reveals Drug Targets and Potential Drug-Repurposing. bioRxiv. 2020; 2020.03.22.002386. doi:10.1101/2020.03.22.002386

12. Keating GM. Sofosbuvir: a review of its use in patients with chronic hepatitis C. Drugs. 2014;74: 1127–1146. doi:10.1007/s40265-014-0247-z

13. de Freitas CS, Higa LM, Sacramento CQ, Ferreira AC, Reis PA, Delvecchio R, et al. Yellow fever virus is susceptible to sofosbuvir both in vitro and in vivo. PLoS Negl Trop Dis. 2019;13: e0007072. doi:10.1371/journal.pntd.0007072

14. Ferreira AC, Reis PA, de Freitas CS, Sacramento CQ, Villas Boas Hoelz L, Bastos MM, et al. Beyond members of the Flaviviridae family, sofosbuvir also inhibits chikungunya virus replication. Antimicrob Agents Chemother. 2018. doi:10.1128/aac.01389-18

15. Ferreira AC, Zaverucha-do-Valle C, Reis PA, Barbosa-Lima G, Vieira YR, Mattos M, et al. Sofosbuvir protects Zika virus-infected mice from mortality, preventing short- and long-term sequelae. Scientific Reports. 2017;7: 9409. doi:doi:10.1038/s41598-017-09797-8

16. Sacramento CQ, de Melo GR, de Freitas CS, Rocha N, Hoelz LV, Miranda M, et al. The clinically approved antiviral drug sofosbuvir inhibits Zika virus replication. Sci Rep. 2017;7: 40920. doi:10.1038/srep40920

17. SOVALDI (sofosbuvir). : 37.

18. Tissue expression of CTSA - Staining in lung - The Human Protein Atlas. [cited 15 Jun 2020]. Available: https://www.proteinatlas.org/ENSG00000064601-CTSA/tissue/lung

19. Tissue expression of CES1 - Summary - The Human Protein Atlas. [cited 15 Jun 2020]. Available: https://www.proteinatlas.org/ENSG00000198848-CES1/tissue

20. Tissue expression of HINT1 - Summary - The Human Protein Atlas. [cited 15 Jun 2020]. Available: https://www.proteinatlas.org/ENSG00000169567-HINT1/tissue

21. Gao Y, Yan L, Huang Y, Liu F, Zhao Y, Cao L, et al. Structure of the RNA-dependent RNA polymerase from COVID-19 virus. Science. 2020;368: 779–782. doi:10.1126/science.abb7498

22. Gordon CJ, Tchesnokov EP, Woolner E, Perry JK, Feng JY, Porter DP, et al. Remdesivir is a direct-acting antiviral that inhibits RNA-dependent RNA polymerase from severe acute respiratory syndrome coronavirus 2 with high potency. J Biol Chem. 2020; jbc.RA120.013679. doi:10.1074/jbc.RA120.013679

23. Ju J, Kumar S, Li X, Jockusch S, Russo JJ. Nucleotide Analogues as Inhibitors of Viral Polymerases. bioRxiv. 2020; 2020.01.30.927574. doi:10.1101/2020.01.30.927574

24. Mesci P, Macia A, Saleh A, Martin-Sancho L, Yin X, Snethlage C, et al. Sofosbuvir protects human brain organoids against SARS-CoV-2. bioRxiv. 2020; 2020.05.30.125856. doi:10.1101/2020.05.30.125856

25. Asadi-Pooya AA, Simani L. Central nervous system manifestations of COVID-19: A systematic review. J Neurol Sci. 2020;413: 116832. doi:10.1016/j.jns.2020.116832

26. Zhou F, Yu T, Du R, Fan G, Liu Y, Liu Z, et al. Clinical course and risk factors for mortality of adult inpatients with COVID-19 in Wuhan, China: a retrospective cohort study. The Lancet. 2020;395: 1054–1062. doi:10.1016/S0140-6736(20)30566-3

27. Thomsen R, Christensen MH. MolDock: a new technique for high-accuracy molecular docking. J Med Chem. 2006;49: 3315–3321. doi:10.1021/jm051197e

28. Rangan R, Zheludev IN, Hagey RJ, Pham EA, Wayment-Steele HK, Glenn JS, et al. RNA genome conservation and secondary structure in SARS-CoV-2 and SARS-related viruses: a first look. RNA. 2020;26: 937–959. doi:10.1261/rna.076141.120

29. Arshad U, Pertinez H, Box H, Tatham L, Rajoli RKR, Curley P, et al. Prioritization of Anti-SARS-Cov-2 Drug Repurposing Opportunities Based on Plasma and Target Site Concentrations Derived from their Established Human Pharmacokinetics. Clin Pharmacol Ther. 2020. doi:10.1002/cpt.1909

30. Zeitlinger M, Koch BCP, Bruggemann R, De Cock P, Felton T, Hites M, et al. Pharmacokinetics/Pharmacodynamics of Antiviral Agents Used to Treat SARS-CoV-2 and Their Potential Interaction with Drugs and Other Supportive Measures: A Comprehensive Review by the PK/PD of Anti-Infectives Study Group of the European Society of Antimicrobial Agents. Clin Pharmacokinet. 2020. doi:10.1007/s40262-020-00924-9

31. Venisse N, Peytavin G, Bouchet S, Gagnieu M-C, Garraffo R, Guilhaumou R, et al. Concerns about pharmacokinetic (PK) and pharmacokinetic-pharmacodynamic (PK-PD) studies in the new therapeutic area of COVID-19 infection. Antiviral Res. 2020; 104866. doi:10.1016/j.antiviral.2020.104866

32. Alexander SPH, Armstrong JF, Davenport AP, Davies JA, Faccenda E, Harding SD, et al. A rational roadmap for SARS-CoV-2/COVID-19 pharmacotherapeutic research and development: IUPHAR Review 29. Br J Pharmacol. 2020. doi:10.1111/bph.15094

33. Dong E, Du H, Gardner L. An interactive web-based dashboard to track COVID-19 in real time. The Lancet Infectious Diseases. 2020;0. doi:10.1016/S1473-3099(20)30120-1

34. Lin L, Lu L, Cao W, Li T. Hypothesis for potential pathogenesis of SARS-CoV-2 infection-a review of immune changes in patients with viral pneumonia. Emerg Microbes Infect. 2020;9: 727–732. doi:10.1080/22221751.2020.1746199

35. Li H, Liu L, Zhang D, Xu J, Dai H, Tang N, et al. SARS-CoV-2 and viral sepsis: observations and hypotheses. The Lancet. 2020;395: 1517–1520. doi:10.1016/S0140-6736(20)30920-X

36. Wang Y, Liu S, Liu H, Li W, Lin F, Jiang L, et al. SARS-CoV-2 infection of the liver directly contributes to hepatic impairment in patients with COVID-19. J Hepatol. 2020. doi:10.1016/j.jhep.2020.05.002

37. Harrison C. Coronavirus puts drug repurposing on the fast track. Nat Biotechnol. 2020. doi:10.1038/d41587-020-00003-1

38. Figueiredo-Mello C, Casadio LVB, Avelino-Silva VI, Yeh-Li H, Sztajnbok J, Joelsons D, et al. Efficacy of sofosbuvir as treatment for yellow fever: protocol for a randomised controlled trial in Brazil (SOFFA study). BMJ Open. 2019;9: e027207. doi:10.1136/bmjopen-2018-027207

39. Pruijssers AJ, George AS, Schäfer A, Leist SR, Gralinksi LE, Dinnon KH, et al. Remdesivir Inhibits SARS-CoV-2 in Human Lung Cells and Chimeric SARS-CoV Expressing the SARS-CoV-2 RNA Polymerase in Mice. Cell Rep. 2020;32: 107940. doi:10.1016/j.celrep.2020.107940

40. Murphy BG, Perron M, Murakami E, Bauer K, Park Y, Eckstrand C, et al. The nucleoside analog GS-441524 strongly inhibits feline infectious peritonitis (FIP) virus in tissue culture and experimental cat infection studies. Vet Microbiol. 2018;219: 226–233. doi:10.1016/j.vetmic.2018.04.026

41. Yan VC, Muller FL. Advantages of the Parent Nucleoside GS-441524 over Remdesivir for Covid-19 Treatment. ACS Med Chem Lett. 2020;11: 1361–1366. doi:10.1021/acsmedchemlett.0c00316

42. HCV drugs sofosbuvir, daclatasvir show promise as potential COVID-19 treatment. [cited 28 Jul 2020]. Available: https://www.healio.com/news/infectious-disease/20200709/hcv-drugs-sofosbuvir-daclatasvir-show-promise-as-potential-covid19-treatment

43. Garcez PP, Loiola EC, Madeiro da Costa R, Higa LM, Trindade P, Delvecchio R, et al. Zika virus impairs growth in human neurospheres and brain organoids. Science. 2016;352: 816–818. doi:10.1126/science.aaf6116

44. CDC. Coronavirus Disease 2019 (COVID-19). In: Centers for Disease Control and Prevention [Internet]. 11 Feb 2020 [cited 30 Mar 2020]. Available: https://www.cdc.gov/coronavirus/2019-ncov/lab/rt-pcr-panel-primer-probes.html

45. Wölfel R, Corman VM, Guggemos W, Seilmaier M, Zange S, Müller MA, et al. Virological assessment of hospitalized patients with COVID-2019. Nature. 2020;581: 465–469. doi:10.1038/s41586-020-2196-x

46. National Health Statistics Reports, Number 122, December 20, 2018. 2018; 16.

47. Bosgra S, van Eijkeren J, Bos P, Zeilmaker M, Slob W. An improved model to predict physiologically based model parameters and their inter-individual variability from anthropometry. Crit Rev Toxicol. 2012;42: 751–767. doi:10.3109/10408444.2012.709225

48. Williams LR. Reference values for total blood volume and cardiac output in humans. Oak Ridge National Lab., TN (United States); 1994 Sep. Report No.: ORNL/TM-12814. doi:10.2172/10186900

49. Sun D, Lennernas H, Welage LS, Barnett JL, Landowski CP, Foster D, et al. Comparison of human duodenum and Caco-2 gene expression profiles for 12,000 gene sequences tags and correlation with permeability of 26 drugs. Pharm Res. 2002;19: 1400–1416. doi:10.1023/a:1020483911355

50. Gertz M, Harrison A, Houston JB, Galetin A. Prediction of human intestinal first-pass metabolism of 25 CYP3A substrates from in vitro clearance and permeability data. Drug Metab Dispos. 2010;38: 1147–1158. doi:10.1124/dmd.110.032649

51. Rodgers T, Rowland M. Physiologically based pharmacokinetic modelling 2: predicting the tissue distribution of acids, very weak bases, neutrals and zwitterions. J Pharm Sci. 2006;95: 1238–1257. doi:10.1002/jps.20502

52. Daclatasvir. [cited 28 Jul 2020]. Available: https://www.drugbank.ca/drugs/DB09102

53. Wang Q, Li W, Zheng M, Eley T, LaCreta F, Garimella T. Physiologically-Based Simulation of Daclatasvir Pharmacokinetics With Antiretroviral Inducers and Inhibitors of Cytochrome P450 and Drug Transporters. : 14.

