## Supplementary material for "The *in vitro* antiviral activity of the anti-hepatitis C virus (HCV) drugs daclatasvir and sofosbuvir against SARS-CoV-2": SARS-CoV-2 and SOF-DAC_Updated SI

Running-title: SARS-CoV-2 susceptibility to daclatasvir and sofosbuvir *in vitro*

Carolina Q. Sacramento[1,10#], Natalia Fintelman-Rodrigues[1, 10#], Jairo R. Temerozo[2,3] , Aline de Paula Dias Da Silva[1,10] Suelen da Silva Gomes Dias[1], André C. Ferreira[1,4,10], Mayara Mattos[1,10], Camila R. R. Pão[1], Caroline S. de Freitas[1, 10], Vinicius Cardoso Soares[1], Lucas Villas Bôas Hoelz[5], Tácio Vinício Amorim Fernandes[5,6], Frederico Silva Castelo Branco[5], Mônica Macedo Bastos[5], Núbia Boechat[5], Felipe B. Saraiva[7], Marcelo Alves Ferreira[7,12], Rajith K. R. Rajoli[8], Carolina S. G. Pedrosa[9], Gabriela Vitória[9], Letícia R. Q. Souza[9], Livia Goto-Silva[9], Marilia Zaluar Guimarães[9,10], Stevens K. Rehen[9,10], Andrew Owen[8], Fernando A. Bozza[9,11], Dumith Chequer Bou-Habib[2,3], Patrícia T. Bozza[1], Thiago Moreno L. Souza[1,12,*]

### - These authors contributed equally to this work

1 – Laboratório de Imunofarmacologia, Instituto Oswaldo Cruz (IOC), Fundação Oswaldo Cruz (Fiocruz), Rio de Janeiro, RJ, Brazil.

2 – National Institute for Science and Technology on Neuroimmunomodulation (INCT/NIM), IOC, Fiocruz, Rio de Janeiro, RJ, Brazil.

3 – Laboratório de Pesquisas sobre o Timo, IOC, Fiocruz, Rio de Janeiro, RJ, Brazil.

4 - Universidade Iguaçu, Nova Iguaçu, RJ, Brazil.

5 – Instituto de Tecnologia de Fármacos (Farmanguinhos), Fiocruz, Rio de Janeiro, RJ, Brazil.

6 – Laboratório de Macromoléculas, Diretoria de Metrologia Aplicada às Ciências da Vida, Instituto Nacional de Metrologia, Qualidade e Tecnologia - INMETRO, Duque de Caxias, RJ 25250-020, Brazil

7 - Instituto de Tecnologia em Imunobiológicos (Bio-Manguinhos), Fiocruz, Rio de Janeiro, RJ, Brazil

8 - Department of Molecular and Clinical Pharmacology, University of Liverpool, Liverpool, L7 3NY, UK;

9 – Instituto D'Or de Pesquisa e Ensino, Rio de Janeiro, RJ, Brazil

10 - Instituto de Ciências Biomédicas, Universidade Federal do Rio de Janeiro, Rio de Janeiro, RJ, Brazil.

11 - Instituto Nacional de Infectologia Evandro Chagas, Fiocruz, Rio de Janeiro, RJ, Brazil

12 - National Institute for Science and Technology on Innovation in Diseases of Neglected Populations (INCT/IDPN), Center for Technological Development in Health (CDTS), Fiocruz, Rio de Janeiro, RJ, Brazil.

### SUPPORTING INFORMATION

***Correspondence footnote:**

Thiago Moreno L. Souza, PhD

*************************************

Fundação Oswaldo Cruz (Fiocruz)

Centro de Desenvolvimento Tecnológico em Saúde (CDTS)

Instituto Oswaldo Cruz (IOC)

Pavilhão Osório de Almeida, sala 16

Av. Brasil 4365, Manguinhos, Rio de Janeiro - RJ, Brasil, CEP 21060340

### SUPPORTING INFORMATION

Figure S1

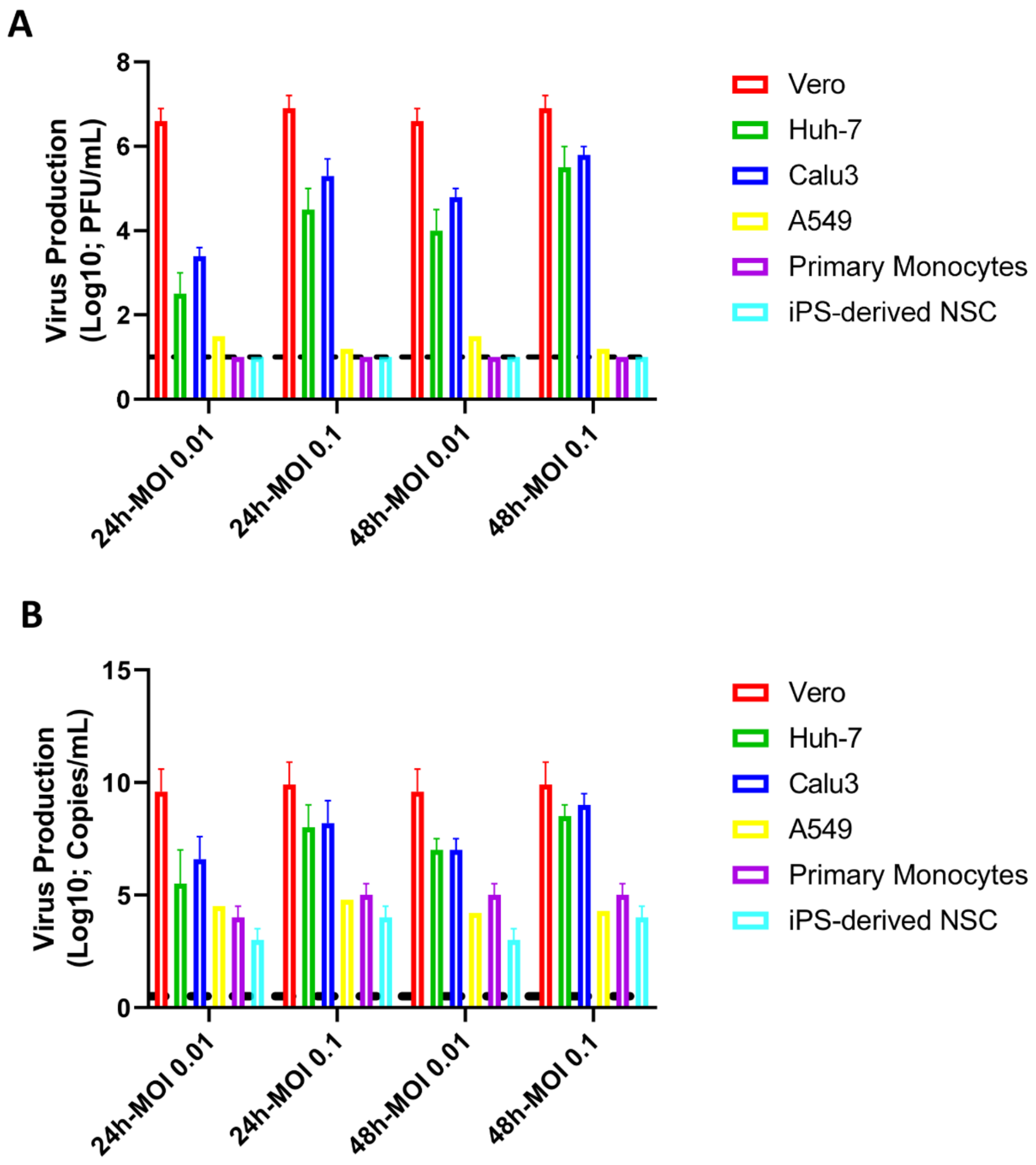


**Figure S1 – SARS-CoV-2 production in different cells lines.** The indicated cell lines, at density of 5 x $10^5$ cells/well in 48-well plates were infected for 1h at 37 °C at MOI of 0.1 or 0.01. Inoculum was removed, cells were washed and incubated with fresh DMEM containing 2% fetal bovine serum (FBS). After 24 or 48 h post-infection, supernatants were collect for titration by plaque forming units (PFU) in Vero cells (A) or by RT-PCR (B). The dashed line indicates the limit of detection. Data represent means ± S.E.M. of three independent experiments.

Figure S2

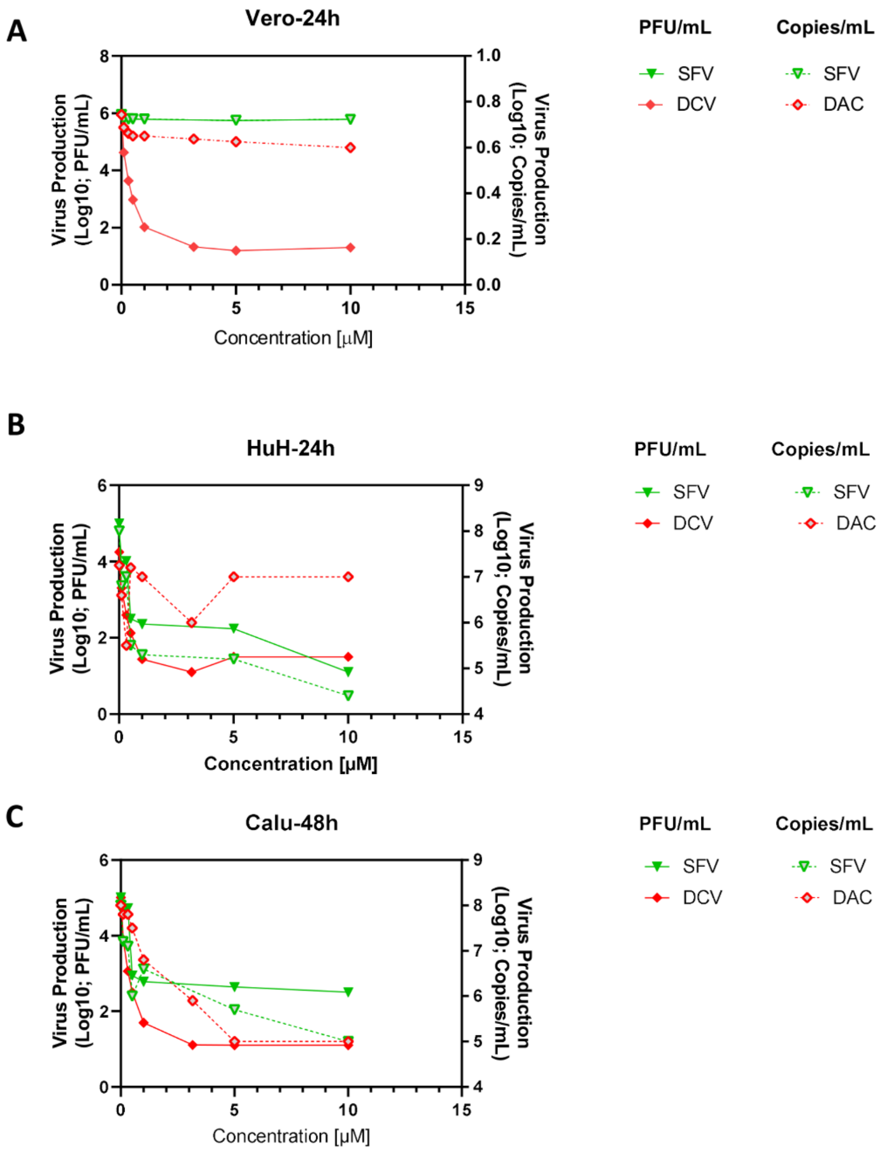


**Figure S2. The antiviral activity of daclatasvir (DCV) and sofosbuvir (SFV) against SARS-CoV-2.** Vero (A), HuH-7 (B) or Calu-3 (C) cells, at density of 5 x $10^5$ cells/well in 48-well plates, were infected with SARS-CoV-2, for 1h at 37 °C. Inoculum was removed, cells were washed and incubated with fresh DMEM containing 2% fetal bovine serum (FBS) and the indicated concentrations of the DCV and SFV. Vero cells (A) were infected with MOI of 0.01 and supernatants were accessed after 24 h. HuH-7 (B) and Calu-3 (C) cells were infected with MOI of 0.1 and supernatants were accessed after 48 h. Viral replication in the culture supernatant was measured by PFU/mL or virus RNA levels. The data represent means ± SEM of three independent experiments.

#### SUPPORTING INFORMATION

Figure S3

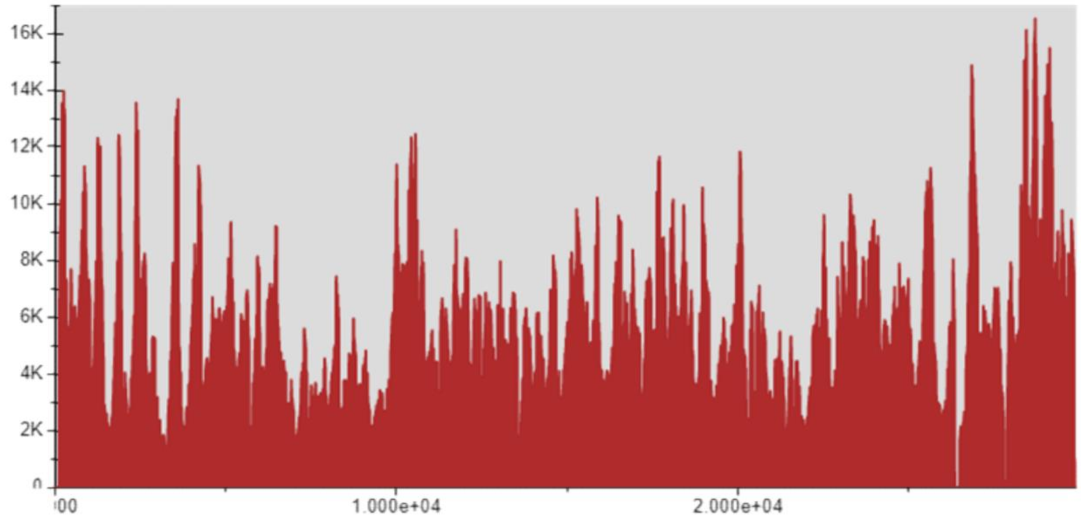


**Figure S3 – Coverage of the SARS-CoV-2 genome.** The representative graphic demonstrate the depth of the SARS-CoV-2 RNA sequence using MGI-2000. X-axis represents virus nucleotides. Y-axis represents the number of reads/site.

### SUPPORTING INFORMATION

Figure S4

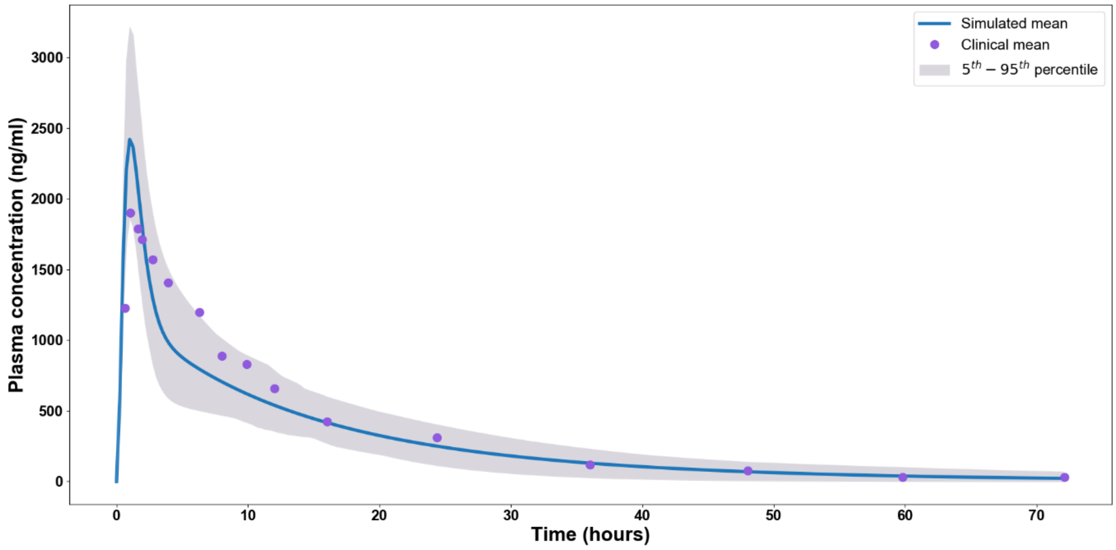


**Figure S4** Daclatasvir PBPK model validation against clinical data [2] for a 100 mg single dose

Figure S5

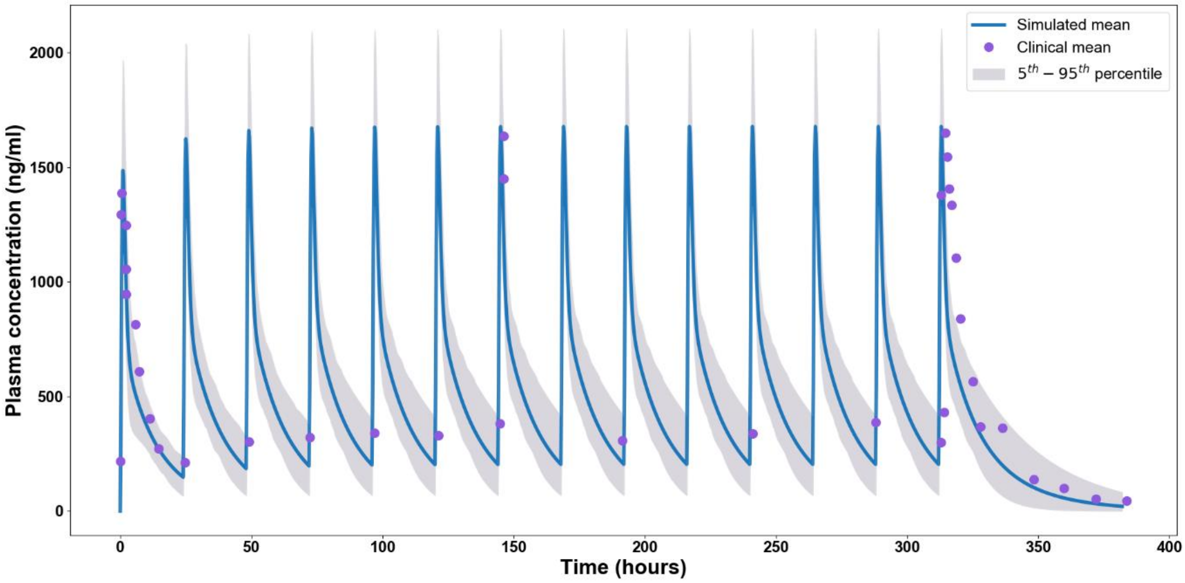


**Figure S5** Daclatasvir PBPK model validation against clinical data [2] for multiple 60 mg OD doses

Figure S6

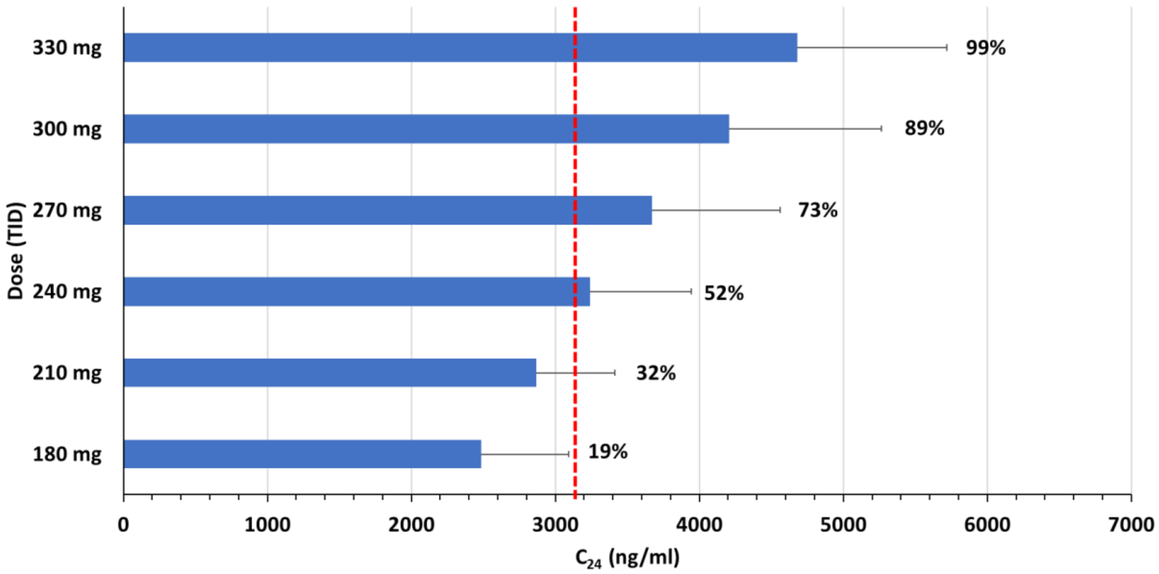


**Figure S6** Daclatasvir PBPK model predictions of the trough plasma concentration at 24 h time point for various TID doses. The red line represents the $EC_{90}$ value (3079 ng/ml) and the percentages adjacent to each of the bars represent the percent of simulated population over $EC_{90}$.

Figure S7

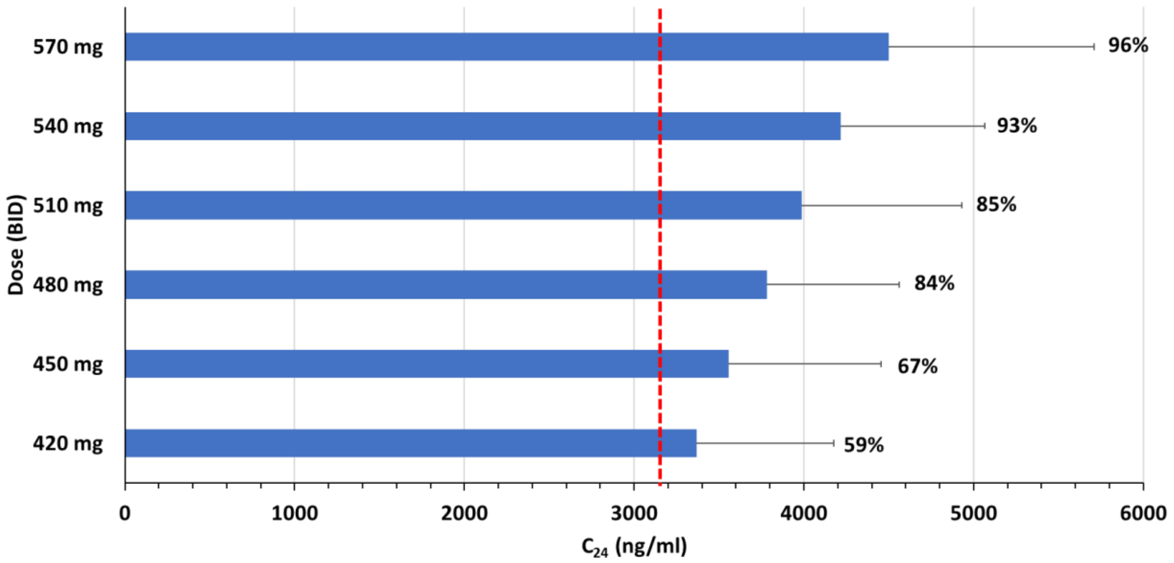


**Figure S7** Daclatasvir PBPK model predictions of the trough plasma concentration at 24 h time point for various BID doses. The red line represents the $EC_{90}$ value (3079 ng/ml) and the percentages adjacent to each of the bars represent the percent of simulated population over $EC_{90}$.

### SUPPORTING INFORMATION

**Table S1** Daclatasvir PBPK input parameters

| Parameter | Daclatasvir |
|---|---|
| Molecular weight | 738.89 [1] |
| Protein binding | 99.4% [2] |
| Log P | 4.05 [2] |
| pKa (diprotic base) | 5.6, 4.9 [2] |
| Blood-to-plasma ratio | 0.8 [2] |
| Apparent permeability, PAMPA (cm/s) | 49e-6 [2] |
| Systemic clearance (L/h) | 4.2 [1] |
| Volume of distribution (L) | 47 [1] |
| Tissue to plasma ratio factor | 0.025 |
| Half-life (h) | 12-15 [1] |

**Table S2** Daclatasvir validation for various single doses [2]

| Dose (mg) | $C_{max}$ (ng/ml) | | | $AUC_{last}$ (ng.h/ml) | | | $C_{24}$ (ng/ml) |
|---|---|---|---|---|---|---|---|
| | Observed | Simulated | Sim/Obs. | Observed | Simulated | Sim/Obs. | Simulated |
| **1** | 16 | 24 ± 4 | 1.50 | 160 | 203 ± 35 | 1.27 | 2.0 ± 1.0 |
| **10** | 200 | 244 ± 46 | 1.22 | 2053 | 2020 ± 452 | 0.98 | 24 ± 10 |
| **25** | 406 | 616 ± 126 | 1.52 | 3962 | 5081 ± 1084 | 1.28 | 60 ± 24 |
| **50** | 1226 | 1192 ± 206 | 0.97 | 13255 | 7999 ± 1699 | 0.60 | 122 ± 43 |
| **100** | 1921 | 2453 ± 443 | 1.28 | 22241 | 16065 ± 2987 | 0.72 | 249 ± 84 |
| **200** | 2816 | 4945 ± 899 | 1.76 | 31473 | 32999 ± 6493 | 1.05 | 473 ± 193 |

### SUPPORTING INFORMATION

**Table S3** Daclatasvir validation for various multiple OD doses [8]

| | $C_{max}$ (ng/ml) | | | $AUC_{last\text{-}dose}$ (ng.h/ml) | | | $C_{24}$ (ng/ml) |
|---|---|---|---|---|---|---|---|
| **Dose (mg)** | **Observed** | **Simulated** | **Sim/Obs.** | **Observed** | **Simulated** | **Sim/Obs.** | **Simulated** |
| **1** | 16 | 28 ± 4 | 1.75 | 125 | 209 ± 46 | 1.67 | 3.0 ± 1.0 |
| **10** | 257 | 276 ± 42 | 1.07 | 2454 | 2108 ± 487 | 0.86 | 25 ± 9.0 |
| **30** | 734 | 847 ± 138 | 1.15 | 6275 | 6303 ± 1459 | 1.00 | 70 ± 30 |
| **60** | 1582 | 1622 ± 260 | 1.03 | 15666 | 12579 ± 2932 | 0.80 | 151 ± 56 |

### SUPPORTING INFORMATION

#### References


1. DrugBank. *Daclatasvir*. 2020 [cited 2020 25/06/2020]; Available from: https://www.drugbank.ca/drugs/DB09102.

2. Qi Wang, W.L., Ming Zheng, Timothy Eley, Frank LaCreta, Tushar Garimella. *Physiologically-Based Simulation of Daclatasvir Pharmacokinetics With Antiretroviral Inducers and Inhibitors of Cytochrome P450 and Drug Transporters*. in *17th International Workshop on Clinical Pharmacology of HIV & Hepatitis Therapy*. 2016. Washington, DC, Available from: http://regist2.virology-education.com/2016/17HIVHEPPK/37_Eley.pdf.
